# CXCL13-CXCR5 Signaling in CD8⁺ T Cell Recruitment and Lymphoid Immune Organization in Clear Cell Renal Cell Carcinoma

**DOI:** 10.64898/2026.05.04.722710

**Authors:** Daniel D. Shapiro, Kye D. Nichols, Moon Hee Lee, Pavlos Msaouel, Yuanshan Li, Yang Zong, Rong Hu, Wei Huang, Karla Esbona, Toshi Kinoshita, Paz Lotan, Daniel F. Roadman, Everlyne Nkadori, Stephanie M. McGregor, David J. Beebe, Sheena C. Kerr, Christian M. Capitini, E. Jason Abel

## Abstract

Clear cell renal cell carcinoma (ccRCC) exhibits heterogeneity in immune infiltration and clinical outcomes, but the mechanisms governing recruitment and organization of tumor-reactive CD8⁺ T cells remain incompletely defined. We investigated the role of the CXCL13–CXCR5 axis in shaping CD8⁺ T cell recruitment, differentiation, and immune organization in high-risk, non-metastatic ccRCC. Human tumor, plasma, and matched adjacent kidney specimens were analyzed using ELISA, quantitative PCR, migration assays, multiplex immunofluorescence, single-cell RNA sequencing, spatial transcriptomics, and a syngeneic mouse model. CXCL13 was among the most upregulated chemokines in ccRCC relative to matched normal kidney and was embedded within a CD8⁺ T cell-associated inflammatory transcriptional program. In transwell and microphysiological system (MPS) assays, CXCL13 promoted CD8⁺ T cell migration, enriched CXCR5⁺ cells among migrating CD8⁺ T cells and showed reduced migration after CXCL13 or CXCR5 blockade. Single-cell analyses identified *CXCR5* expression within stem-like CD8⁺ T cell states associated with *TCF7* and *IL7R*, whereas *CXCL13* associated with later cytotoxic/exhausted states along a continuous differentiation landscape. Spatial transcriptomics demonstrated that stem-like CD8⁺ T cells localized within structured lymphoid aggregates enriched for B cells, coordinated CXCL13/CXCR5 expression, and signaling programs. *In vivo*, tumor-derived CXCL13 suppressed tumor growth, increased intratumoral CD8⁺ T cell infiltration, and enriched CXCR5⁺TCF1⁺CD8⁺ stem-like T cells. In human tumors, higher CXCL13 expression correlated with increased CXCR5⁺CD8⁺ T cell infiltration and improved recurrence-free survival. These findings identify CXCL13 as a regulator of immune recruitment and niche organization and support the CXCL13–CXCR5 axis as a biomarker and possible therapeutic target in ccRCC.

## INTRODUCTION

Clear cell renal cell carcinoma (ccRCC) is an immunogenic malignancy characterized by marked variability in clinical outcomes [1–4]. Patients with non-metastatic disease and adverse pathologic features such as larger tumors, advanced T stage, and higher nuclear grade remain at substantial risk for progression to metastatic disease following surgery, yet current clinicopathologic features incompletely predict this variability in outcomes [5]. Although intratumoral CD8⁺ T cell infiltration has been associated with improved outcomes in non-metastatic disease [1, 6, 7], some patients with T cell-inflamed tumors ultimately develop metastatic disease, suggesting that functional and spatial heterogeneity within the CD8⁺ T cell compartment critically influences anti-tumor immunity [8].

The classical paradigm of anti-tumor immunity proposes that tumor-specific T cells are primed in tumor-draining lymph nodes, then migrate into tumors and exert cytotoxic activity [9, 10]. However, emerging data identify a distinct population of intratumoral CD8⁺ T cells with stem-like properties that sustain the anti-tumor response [7, 11]. These “stem-like” or “progenitor” cells, characterized by expression of TCF1 (*TCF7*), retain proliferative capacity and may give rise to more differentiated effector and exhausted T cell populations, thereby maintaining the intratumoral T cell response and contributing to durable tumor control and improved immunotherapy response [11–14].

Despite increasing recognition of this population, a fundamental gap remains in understanding how stem-like CD8⁺ T cells are recruited to and spatially organized within the tumor microenvironment. Prior work suggests that these cells localize within specialized intratumoral niches, often associated with antigen-presenting cells, vascular structures, or tertiary lymphoid structure (TLS)-like regions, where local signals support their maintenance and differentiation [7, 14–16]. However, the mechanisms governing their trafficking into tumors remain incompletely defined and these findings require validation.

The chemokine receptor CXCR5 has emerged as another potential marker of stem-like CD8⁺ T cells across chronic infection and cancer [12, 13, 17]. CXCR5⁺CD8⁺ T cells exhibit features of stem-like cells, including expression of TCF7, proliferative capacity, and the ability to differentiate into more terminally exhausted populations [12, 13, 17]. CXCR5 is the canonical receptor for CXCL13, a chemokine classically involved in B cell and T follicular helper cell trafficking and lymphoid organization [18–20]. While CXCL13 has been implicated in lymphoid architecture and anti-tumor immune responses among other cancer types, its role in ccRCC and relationship to CD8⁺ T cell recruitment, differentiation, and spatial organization remain poorly defined [18, 21–24].

We hypothesized that CXCL13 regulates the recruitment and spatial organization of CXCR5⁺CD8⁺ stem-like T cells within the ccRCC tumor microenvironment. To test this, we integrated analyses of human tumor specimens with functional migration assays, single-cell RNA sequencing, spatial transcriptomic profiling, and *in vivo* tumor models to determine how CXCL13-CXCR5 signaling shapes CD8⁺ T cell recruitment, differentiation, and spatial immune organization in ccRCC.

## METHODS

### Clear cell renal cell carcinoma tumor sampling

Following institutional review board approval (#2024-0997), tumor and matched adjacent kidney tissue were obtained from patients with non-metastatic clear cell renal cell carcinoma tumors at high risk of recurrence after surgery (defined as tumors ≥7cm in size). Patients provided informed consent for tissue and peripheral blood collection. Preoperative blood samples were collected for plasma isolation and CD8⁺ T cell extraction.

### Cell lines and reagents

Human RCC cell lines and RENCA murine cells were obtained from commercial sources (ATCC) and maintained under standard culture conditions [25]. CXCL13 overexpression and knockout cell lines were generated using lentiviral constructs (pEZ-mCXCL13-Flag and pLenti-U6-sgRNA-Cas9) followed by antibiotic selection (Genecopoeia; ABM Inc.). Neutralizing anti-CXCL13, anti-CXCR5, control IgG antibodies were purchased from R&D systems. Recombinant human and mouse CXCR13 and CXCL9 protein was purchased from Gibco.

### Tumor-derived epithelial cells and CD8^+^ T cell isolation

Tumor specimens were enzymatically dissociated into single-cell suspensions. Tumor-derived epithelial cells were enriched by negative selection with anti-CD31 microbeads and expanded in culture to generate tumor derived cell cultures. CD8⁺ T cells were isolated from tumor using the REAlease CD8 microbead kit (Miltenyi Biotec) with bead removal and peripheral blood using the MACSxpress negative selection kit, with purity confirmed by flow cytometry.

### Migration assays

For transwell migration assays, 2×10^5^ tumor cells or RCC cell lines were plated in the lower chamber. After 24 hours, media were replaced with serum-free RPMI with or without recombinant human CXCL13 (10nM). CD8^+^ T cells (5×10^4^) were added to upper inserts and allowed to migrate for 2 hours. Migrated cells were collected from the lower chamber and quantified using an automated cell counter (Invitrogen).

Microphysiological system (MPS) lumeNEXT devices were fabricated and assembled as previously described [26–28]. Tumor spheroids were generated using the hanging-drop method with approximately 1,000 epithelial cells for 2 days. Human umbilical vein endothelial cells (HUVEC) cells were seeded to generate endothelial lined lumens one day before spheroid loading. A498 CXCL13 wild-type or knockout spheroids were embedded in collagen hydrogel and introduced into side ports. After 24 hours, devices were rinsed, fixed in 4% PFA, stained with DAPI, and imaged by microscopy.

### CXCL13 quantification and gene expression analysis

CXCL13 concentrations in plasma and tumor lysates were measured using enzyme-linked immunosorbent assays per manufacturer protocols (R&D, DCX130). Gene expression was assessed by quantitative PCR following RNA extraction and reverse transcription using standard methods. Primer sequences are provided in Supplementary Methods.

### Flow cytometry

Single-cell suspensions from tumor were first stained with viability dye eFluor 780 (eBioscience), followed by fixing and permeabilization and staining for CD45 (30-F11), CD3 (500A2), CD8 (53-6.7), TIM3 (RMT3-23), CD4 (RM4-5), CXCR5 (SPRCL5), and CD11b (M1/70) from life technologies, PD1 (NAT105) form Invitrogen, TCF1 (7F11A10) from Biolegend. Samples were analyzed on Attune flow cytometers (ThermoFisher).

### In vivo RCC tumor growth

All animal studies were conducted in accordance with institutional guidelines. Wild-type, CXCL13-overexpressing (“CXCL13 High”), or CXCL13 knockout RENCA cells were injected subcutaneously into the flanks of 6–8-week-old BALB/c mice (n=5 per group). Tumor growth was measured every 3 days for 3 weeks. At endpoint, tumors were harvested, dissociated, and analyzed for immune cell composition.

### Tissue microarray

A tissue microarray (TMA) was constructed from high-risk, non-metastatic ccRCC tumors from patients who did or did not develop metastatic disease after surgery (n=46). Tumors were matched for clinicopathologic features including age, tumor size, grade, and stage. Multiplex immunofluorescence staining for CXCL13, CD8, and CXCR5 was performed as previously described [1, 29]. Slides were imaged and analyzed to quantify CXCL13 optical density and CD8⁺CXCR5⁺ T cell infiltration. Additional details are provided in the Supplementary Methods.

### Multi-modal transcriptomic analysis

To identify biological processes associated with *CXCL13* and *CXCL9*, co-expression modules were constructed from TCGA M0 ccRCC data using weighted gene co-expression network analysis (WGCNA). The soft-thresholding power was selected based on scale-free topology fit and mean network connectivity, with β = 6 chosen for downstream analysis (**Fig S1**). Modules containing *CXCL13* and *CXCL9* were selected and analyzed by overrepresentation analysis using C5 gene sets and Fisher’s exact test.

Publicly available single-cell RNA-seq datasets (Li et al. [30] and Krishna et al. [31]) were analyzed to define CD8⁺ T cell states and transcriptional programs associated with *CXCL13* and *CXCR5*. Trajectory inference was performed using Monocle3 [32], with stem-like CD8⁺ T cells designated as the root population.

To identify shared transcriptional programs across CD8⁺ T cell states, we applied consensus non-negative matrix factorization (cNMF), selecting 13 factors based on reconstruction error (**Fig S2**). To determine genes that may be part of the same cellular programs as CXCL13 and CXCR5, we ranked genes by similarity of their coordinated cNMF program contributions to CXCL13 and CXCR5. We then characterized the two lists of CXCL13- and CXCR5-covarying genes by gene set enrichment using canonical CD8⁺ T cell phenotype gene sets (**Table S1**). Similarly, pairwise gene correlations among cNMF-derived metaprogram genes were used to assess relationships between *CXCR5*, *CXCL13*, and canonical CD8⁺ T cell phenotype genes, with validation by MAGIC, Kendall correlation, and zero-inflated negative binomial modeling.

Visium HD spatial transcriptomic data were analyzed to infer single-cell-level spatial composition and immune organization. CD8⁺ T cell states were quantified using a stem-to-exhausted “transition score” mapped across CD8^+^ T cell annotated pseudocells. Immune aggregates were identified by density-based spatial clustering to delineate spatially organized immune neighborhoods. Aggregate composition was compared using empirical cumulative fractions of continuous CD8⁺ T cell states as well as proportions and counts of immune cell types. Comparative gene expression between cells classified as “lymphoid” aggregates and surrounding immune cells in no-progression slides (largely composed of “stimulated” aggregates) was modeled using scVI. Individual cells were used as observations, with immune cell type included as a covariate and slide number treated as a batch variable. Genes with the highest probability of differential expression were subjected to overrepresentation analysis using Enrichr and C2 Reactome gene sets. Additional details of transcriptomic analyses can be found in Supplementary Methods.

### Statistical Analysis

Statistical analyses were performed using nonparametric tests unless otherwise specified. Group comparisons were conducted using Mann–Whitney U or paired tests as appropriate. Survival analyses were performed using Kaplan–Meier methods with log-rank testing. A two-sided p-value <0.05 was considered statistically significant.

## RESULTS

### CXCL13 is elevated in high-risk non-metastatic ccRCC and co-expressed within a CD8⁺ T cell–associated transcriptional program

We initially evaluated CXCL13 expression compared to other chemokines associated with CD8^+^ T cell migration in tumor compared to matched normal kidney tissue from patients with high-risk, non-metastatic ccRCC. Interestingly, *CXCL13* and *CXCL9* were among the most upregulated chemokine genes in tumors (**Fig. 1A**). While CXCL9 is a well-established mediator of CD8⁺ T cell recruitment, the role of CXCL13 in CD8^+^ T cell recruitment is less defined. Plasma CXCL13 protein concentrations measured by ELISA were higher in patients with ccRCC tumors compared with age-matched healthy controls (**Fig. 1B**), and paired analysis confirmed increased *CXCL13* transcript expression in tumors compared to matched tumor-adjacent normal kidney (**Fig. 1C**). *CXCL13* expression varied substantially across individual tumors (**Fig. 1D**), indicating heterogeneity in CXCL13-associated signaling.

**Figure 1.**
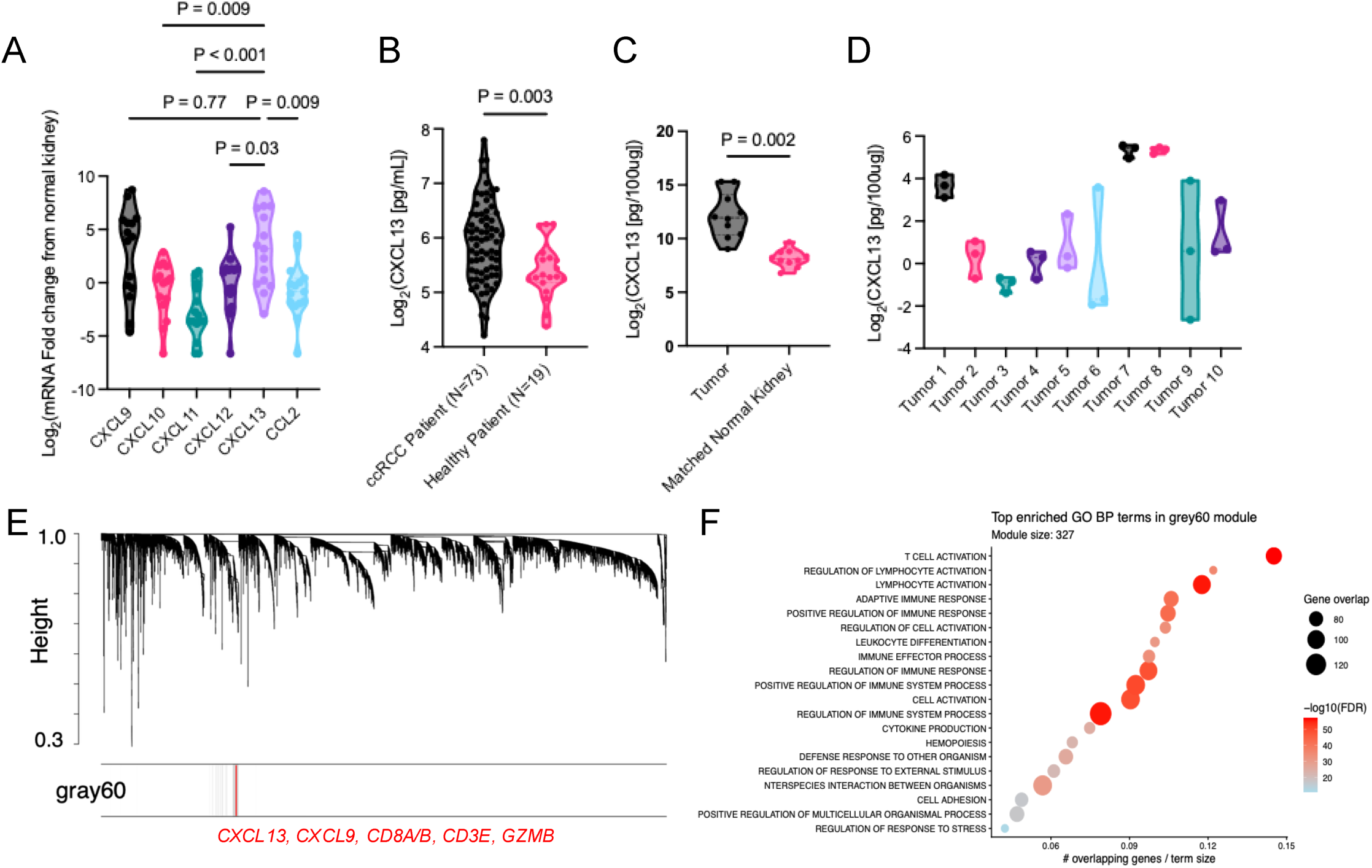
CXCL13 is elevated in ccRCC and co-expressed within a CD8⁺ T cell–associated transcriptional program. (A) Differential expression of selected chemokines in clear cell renal cell carcinoma (ccRCC) tumors compared with matched normal kidney tissue from patients undergoing nephrectomy for non-metastatic ccRCC with tumors >7 cm (Mann–Whitney test; N = 15). (B) Plasma CXCL13 protein concentrations measured by ELISA in patients with ccRCC (N=73) compared with age-matched healthy controls (N=19) (Mann–Whitney test). (C) CXCL13 transcript expression in ccRCC tumors compared with matched normal kidney tissue from the same patients (N=10, Wilcoxon matched-pairs signed-rank test). (D) CXCL13 expression across individual ccRCC tumors, illustrating inter- and intra-tumor heterogeneity. Three spatially separated locations were analyzed from each tumor to evaluate heterogeneity. (E) Dendrogram based on topological overlap similarity generated from Weighted gene co-expression network analysis (WGCNA) showing hierarchical clustering of genes, where x-axis labels represent individual genes and branch height (y-axis) reflects dissimilarity (highest dissimilarity=1.0) between genes and their assigned modules. The highlighted ‘gray60’ module includes *CXCL9, CD8A, CD8B, CD3E, GZMB,* and *CXCL13*. (F) Bubble plot of the top 20 overrepresented Gene Ontology (GO) Biological Process (C5) terms in the grey60 WGCNA module. The x-axis shows the proportion of ‘gray60’ module genes overlapping each GO term. Dot size indicates the number of overlapping genes, and color represents −log10(FDR) from Fisher’s exact test with Benjamini–Hochberg correction. The subtitle reports the total number of genes in the grey60 module.

Furthermore, WGCNA performed on the KIRC TCGA M0 ccRCC data identified a “gray60” co-expression module enriched for *CXCL9, CXCL13*, and canonical CD8⁺ T cell–associated genes (e.g., *CD8A, CD8B, CD3E, GZMB*), in which *CXCL13* was prominently embedded (**Fig. 1E, Table S2**). Gene ontology over-representation analysis further revealed enrichment of T cell activation and cytotoxic effector pathways within this module (**Fig. 1F, Table S3**), supporting CXCL13 as part of an inferred CD8⁺ T cell cytotoxic effector program.

### CXCL13 promotes CD8⁺ T cell migration toward CXCL13-producing cells

Having established that CXCL13 is elevated in ccRCC tumors and associated with a T cell–response transcriptional program, we next evaluated whether CXCL13 promotes CD8⁺ T cell migration. In transwell assays using peripheral blood–derived CD8⁺ T cells from patients with high-risk, non-metastatic ccRCC tumors, migration was minimal under serum-free conditions and modest with serum, while CXCL9 induced robust migration as expected. CXCL13 similarly increased CD8⁺ T cell migration compared to CXCL9 conditions, supporting a chemotactic role for CXCL13 (**Fig. 2A**).

**Figure 2.**
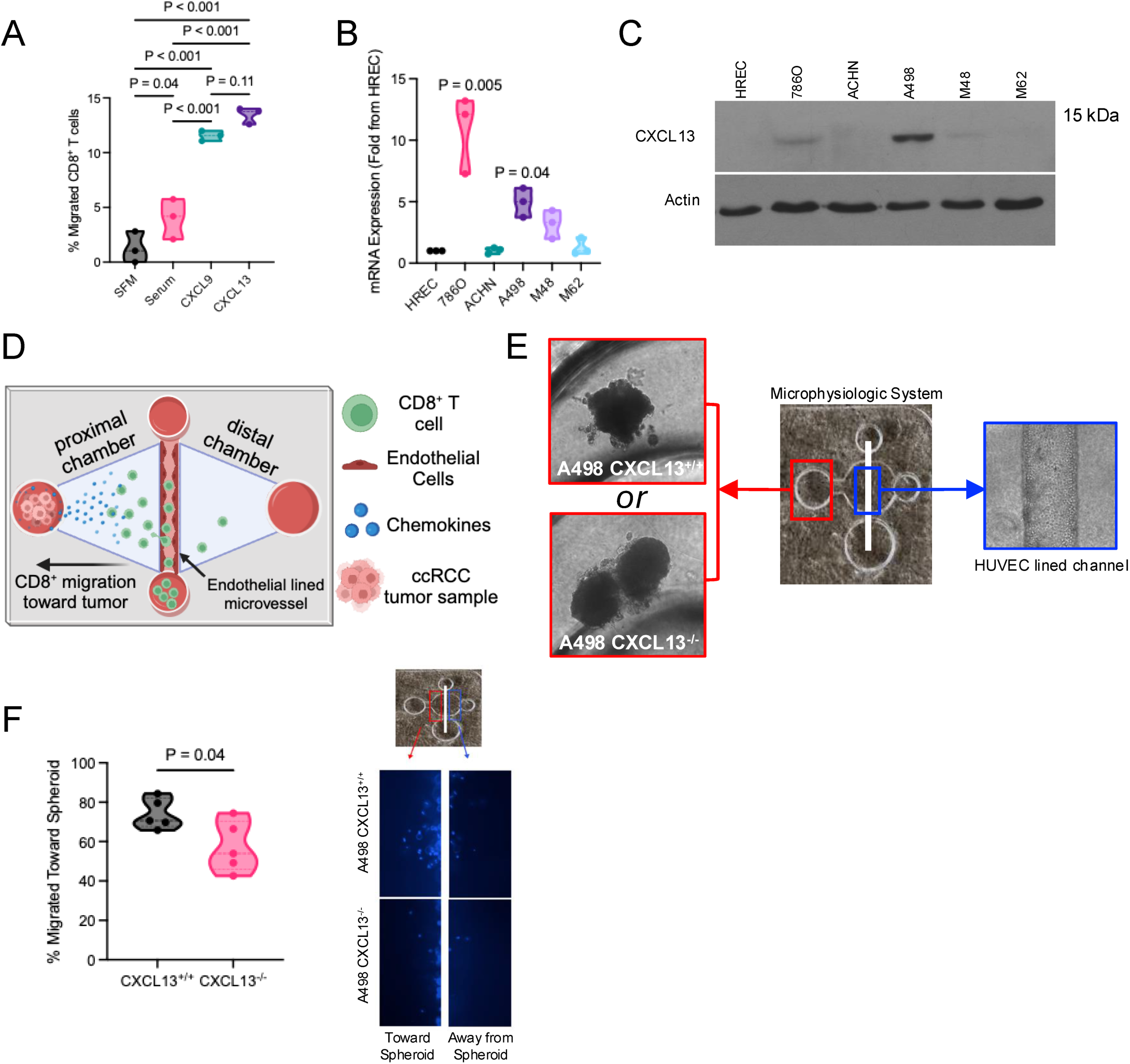
CXCL13 promotes CD8⁺ T cell migration and can be modeled in a microphysiologic ccRCC system. **(A)** Transwell migration assay demonstrating chemotactic responses of CD8⁺ T cells isolated from the peripheral blood of patients with clear cell renal cell carcinoma (ccRCC). Migration was evaluated under serum-free media (SFM), serum-containing media, CXCL9, or CXCL13 stimulation. CXCL9 served as a positive control for CD8⁺ T cell chemotaxis. (N=3, Mann-Whitney test) **(B)** CXCL13 mRNA expression across RCC cell lines and human renal epithelial cells (HREC) measured by quantitative PCR and normalized to HREC expression. Expression compared to HREC (Wilcoxon matched-pairs signed rank test comparing to HREC) **(C)** Western blot analysis demonstrating CXCL13 protein expression across RCC cell lines. **(D)** Schematic of the microphysiologic migration assay. CD8⁺ T cells migrate through an endothelial cell–lined microvessel toward tumor spheroids that produce chemokines. **(E)** Representative images of the microphysiologic system incorporating either CXCL13-expressing A498 tumor spheroids (CXCL13⁺^/^⁺) or CXCL13 knockout A498 spheroids (CXCL13⁻^/^⁻). Tumor spheroids are positioned adjacent to a human umbilical vein endothelial cell (HUVEC)–lined microvascular channel. **(F)** Quantification of CD8⁺ T cell migration toward A498 tumor spheroids in the microphysiologic system (MPS). Representative image showing CD8^+^ T cell migration from endothelial lined channel either toward or away from the A498 tumor spheroid (Wilcoxon matched-pairs signed rank test, N=5).

To further create a model system of CXCL13 expression, we assessed *CXCL13* expression across established RCC cell lines. *CXCL13* mRNA expression varied across lines, with 786O and A498 showing the highest levels relative to HREC (human renal epithelial cells) controls (**Fig. 2B**). CXCL13 protein expression was highest in A498 cells (**Fig. 2C**), which were selected for subsequent assays.

To model chemokine-driven CD8^+^ T cell chemotaxis in a more physiologic context, we utilized a MPS system incorporating A498 tumor spheroids adjacent to an endothelial-lined microvessel (**Fig. 2D**). Using peripheral CD8⁺ T cells from ccRCC patients, migration toward CXCL13-expressing A498 wild-type spheroids was compared with CXCL13 knockout spheroids (**Fig. 2E**). CD8⁺ T cells demonstrated greater migration toward CXCL13-expressing spheroids (**Fig. 2F**), suggesting tumor-derived CXCL13 enhances CD8⁺ T cell trafficking.

### CXCL13 enriches CXCR5⁺ CD8⁺ T cells among migrating lymphocytes and mediates migration through the CXCL13–CXCR5 axis

Having established that CXCL13 promotes CD8⁺ T cell migration, we next sought to determine whether this effect is mediated through its receptor, CXCR5, on CD8⁺ T cells. Flow cytometric analysis of tumor-infiltrating lymphocytes from high-risk, non-metastatic ccRCC tumors confirmed the presence of CXCR5⁺CD8⁺ T cells, with variable frequencies across patients (**Fig. 3A; Fig. S3**). To directly test whether CXCL13 preferentially recruits CXCR5⁺CD8⁺ T cells, we performed transwell assays using CD8⁺ T cells isolated from tumors and quantified CXCR5 expression among migrating cells. Under serum-free conditions, a smaller proportion of migrating CD8⁺ T cells expressed CXCR5, whereas CXCL13 stimulation significantly increased the proportion of CXCR5⁺CD8+ T cells among total migrating CD8⁺ T cells (**Fig. 3B**), indicating selective enrichment of this population.

**Figure 3.**
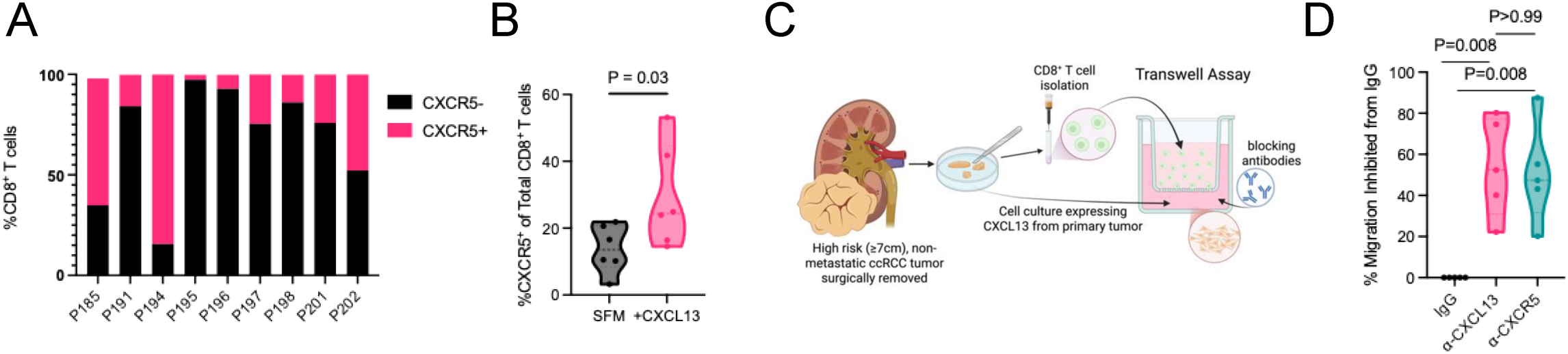
CXCL13 enriches CXCR5⁺CD8⁺ T cells among migrating lymphocytes and migration toward ccRCC tumor cells is mediated through the CXCL13–CXCR5 axis. **(A)** Quantification of CXCR5+CD8+ T cells among total tumor-infiltrating CD8⁺ T cells across individual non-metastatic ccRCC tumors (N=9). **(B)** Percentage of CXCR5⁺ cells among CD8⁺ T cells isolated from ccRCC tumors that migrated in response to serum-free media (SFM) or CXCL13 in transwell assays (Mann–Whitney test, N=6). **(C)** Experimental schematic. Tumor-derived primary ccRCC cell cultures were generated from surgically resected high-risk (≥7cm), non-metastatic tumors. CD8⁺ T cells were isolated and migration toward tumor-derived signals was evaluated using transwell assays in the presence of CXCL13- or CXCR5-blocking antibodies or matched IgG controls. **(D)** Quantification of CD8⁺ T cell migration toward tumor-derived cell cultures in the presence of CXCL13- or CXCR5-blocking antibodies (Wilcoxon matched-pairs signed rank test, N=5).

To confirm that this effect is mediated by tumor microenvironment-derived CXCL13, we generated primary CXCL13-expressing ccRCC cultures from high-risk, non-metastatic tumors and assessed migration of autologous tumor-infiltrating CD8⁺ T cells in transwell assays (**Fig. 3C; Fig. S4**). Because these cultures contained both malignant and non-malignant stromal components, they more closely reflected CXCL13 production within the native tumor microenvironment. Blocking either CXCL13 or its receptor, CXCR5, with neutralizing antibodies reduced CD8⁺ T-cell migration compared with IgG controls (**Fig. 3D**), supporting a role for the CXCL13-CXCR5 axis in mediating trafficking toward primary ccRCC cultures. Together, these findings indicate that CXCL13 promotes migration of CXCR5⁺CD8⁺ T cells.

### scRNA-seq analysis resolves stem-like CD8⁺ T cells within a continuous differentiation landscape in ccRCC characterized by context-dependent CXCR5 and CXCL13 expression

Together, these assays demonstrate that CXCL13 promotes CD8⁺ T cell migration and enriches CXCR5⁺CD8⁺ T cells among migrating lymphocytes. However, these assays do not define the transcriptional identity of the recruited cells or clarify how CXCR5 and CXCL13 relate to CD8⁺ T cell differentiation in ccRCC. We therefore analyzed single-cell RNA-seq datasets to place CXCR5⁺CD8⁺ T cells within the broader CD8⁺ T-cell differentiation landscape. Integration of publicly available single-cell RNA-seq datasets [30, 31] revealed expected immune and tumor cell populations with consistent representation across cohorts (**Fig. 4A–C**). Within CD8⁺ T cells, gene expression previously linked to differentiation and exhaustion showed potential state-specific patterns along a continuum ranging from stem-like to effector, effector memory, and exhausted-like states (**Fig. 4D**) [30].

**Figure 4.**
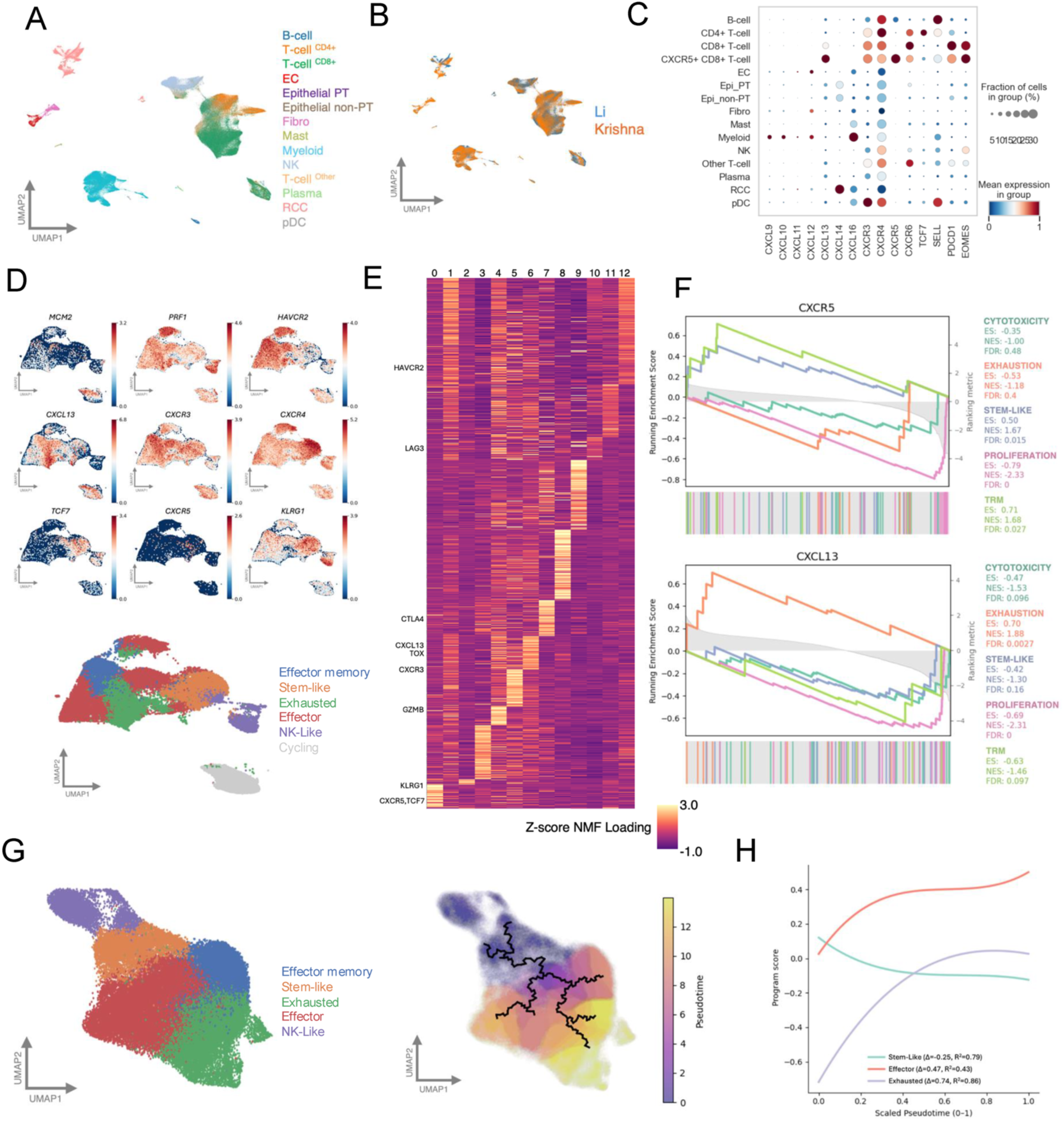
Single-cell transcriptomic analysis identifies CXCR5⁺CD8⁺ T cells as a stem-like population positioned along possible differentiation trajectories in ccRCC. **(A)** 2D UMAP projection of single-cell RNA-seq data derived from Harmony-integrated PCA-reduced feature space, showing the overall cell type composition of the Li et al. and Krishna et al. datasets. Cells are colored by CellTypist-annotated cell type identities. **(B)** 2D UMAP embedding colored by dataset of origin (Li et al. versus Krishna et al.), illustrating strong inter-dataset mixing and concordance following Harmony-based integration. **(C)** Dot plot showing expression of selected chemokine- and immune-related genes across major cell populations and defined CD8⁺ T cell subclusters. Dot color represents scaled (by gene) mean expression, and dot size indicates the percentage of cells expressing each gene within each group. **(D)** UMAP projection representing the landscape of CD8⁺ T cells and colored by specific gene expression relevant to the CD8^+^T cell discrete cell type labels. Cells are grouped by Leiden clustering and annotated using majority-vote, providing a simplified view of the shared CD8⁺ T cell transcriptional landscape across both datasets. **(E)** Heatmap of factorized gene loadings linking genes to transcriptional programs. The plot shows additive gene expression contributions to each metagene program inferred from cNMF. Genes on y axis, hierarchically clustered by loading value show visible patterns and distinct programs. For simplicity, only genes in D are shown. **(F)** A ranked gene set enrichment analysis in which genes are ordered by similarity of their cNMF loading profiles to *CXCL13-* and *CXCR5-associated* programs, quantified using cosine similarity. The enrichment score (ES) reflects the degree to which genes associated with CD8⁺ T cell states (stem-like, cytotoxic, proliferating, and exhausted) are overrepresented in ranked list of genes. Normalized enrichment score and FDR are annotated and colored by gene set. The bottom bars show gene hits that correspond to the ranking shown as shaded gray line. **(G)** UMAP visualization of CD8⁺ T cell states, with an additional UMAP colored by pseudotime values. Monocle-inferred trajectories are overlaid on the latter UMAP to represent the learned principal graph in reduced-dimensional gene-expression space, which defines pseudotime-based ordering of cells from a stem-like root population. **(H)** Line plot where the y-axis shows scores for stem-like, cytotoxic, proliferating, and exhausted gene sets generated using Scanpy’s score_genes function. The x-axis represents Monocle-inferred pseudotime, ordering cells along a trajectory rooted in stem-like CD8⁺ T cells.

Analysis of *CXCR5* and *CXCL13* in the cNMF latent space showed distributed, partially overlapping associations across transcriptional programs, consistent with context-dependent chemokine expression rather than a single biological state (**Fig. 4E**). By ranking genes based on similarity of cNMF loadings, we found CXCR5-associated genes were enriched for stem-like and resident memory programs, whereas CXCL13-associated genes were enriched for exhaustion (**Fig. 4F**). Similarly, gene–gene pairwise correlations alongside MAGIC-imputed expression data demonstrated *CXCR5* to be positively associated with stem-associated genes (*TCF7, IL7R*) and negatively associated with exhaustion-associated genes (*TOX, HAVCR2*) (**Table S4**, **Table S5**).

We next used trajectory inference to assess how *CXCR5* and *CXCL13* expression changed across a differentiation continuum. This identified a continuum from stem-like states toward effector and exhausted states (**Fig. 4G–H**). Gene expression along pseudotime showed *CXCL13* expression increasing along later stages of pseudotime with exhaustion- and effector-associated genes, indicating an association with more differentiated, antigen-experienced CD8⁺ T cell states. Conversely, *CXCR5* expression declined along differentiation trajectories (**Fig. S5**).

We next assessed whether the transcriptional dynamics identified across pseudotime were also preserved at the clonotype level. Using paired αβ TCR sequences to define clonotypes, we found that the stem-like *CXCR5/TCF7* expressing program was maintained across donors and clone structures (**Fig. S6A**). Additionally, expanded clonotypes frequently contained stem-like cells, with 47.7% of expanding clones (≥3 cells) including multiple stem-like cells, and some comprising hundreds of stem-like cells. In contrast, CXCL13 expression was more closely associated with exhaustion-dominant clones characterized by low proportions of stem-like cells (**Fig. S6B**).

### Spatial transcriptomic analysis reveals structured immune niches enriched for stem-like CD8⁺ T cells in ccRCC

To characterize the spatial organization of the ccRCC tumor microenvironment, we performed spatial transcriptomic profiling of tumor sections from patients with high-risk non-metastatic ccRCC who either did (n=4) or did not (n=4) progress to metastatic disease after surgery (**Table S6**). Single-cell estimates from 2μm bin-level data revealed heterogeneous distributions of immune and stromal populations within and across tumors (**Fig. 5A-C; Fig. S7**). Broad cell-type annotations were supported by canonical marker expression and scRNA-seq–derived signatures (**Fig. S8A**). CD8⁺ T cell states (stem-like, effector, and exhausted) clustered across samples reproducibly based on established marker gene expression, with *CXCR5* characterizing the stem-like cluster and *CXCL13* enriching the exhausted cluster (**Fig. S8B**). Stromal area–normalized immune cell counts (density) varied across tumors, with non-progressing tumors showing higher densities of CD8⁺ effector T cells (**Fig. 5D**).

**Figure 5.**
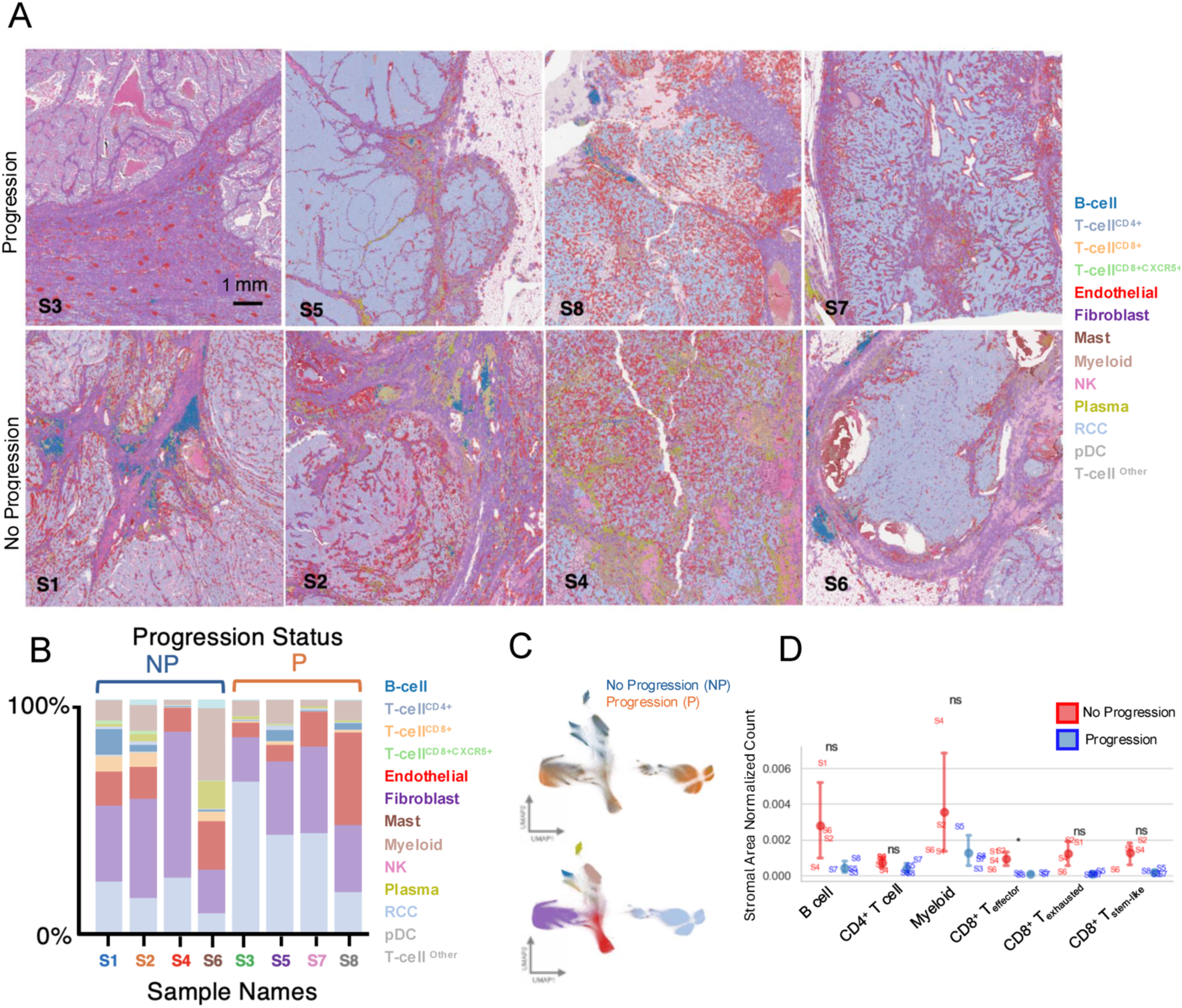
Spatial transcriptomic characterization of the ccRCC tumor microenvironment by progression status. **(A)** H&E overlayed with ENACT-style polygons depicting individual cells. Cells are colored by annotated cell type, as indicated in the legend. Samples are grouped by progression status (progression vs no progression). Scale bar, 1 mm. **(B)**Proportion of annotated cell types across individual samples. Each bar represents a sample, with colors corresponding to cell type annotations shown in (A). Samples are grouped by progression status (NP=no progression, P=progression). **(C)** 2D UMAP projections of scVIVA latent embeddings representing the transcriptomic landscape of pseudocell-estimated expression, colored by progression status (top) and cell type (bottom). **(D)** Quantification of stromal area–normalized cell counts for selected immune cell populations across samples. Points represent individual samples, grouped by progression status. Statistical comparisons were performed using the Mann–Whitney U test (*p<0.05, ns=not significant).

During exploratory spatial analysis, we observed that CD8⁺ T cells, including stem-like subsets, localized preferentially within immune aggregates enriched for B cells and other T cells (**Fig. 6A,B; Fig. S9-S10**). These aggregates also showed co-localization of *CXCR5* and *CXCL13*, supported by Moran’s I spatial autocorrelation of expected expression under the scVIVA model (**Fig. 6C; Fig. S11; Table S7**). In addition, stem-like CD8⁺ T cells and their associated immune aggregates were preferentially located in stromal rather than tumor-rich regions (**Fig. S12**). Together, these findings indicate that stem-like CD8⁺ T cells are concentrated within structured stromal niches enriched for B cells and T cells and characterized by coordinated CXCR5-CXCL13 expression.

**Figure 6.**
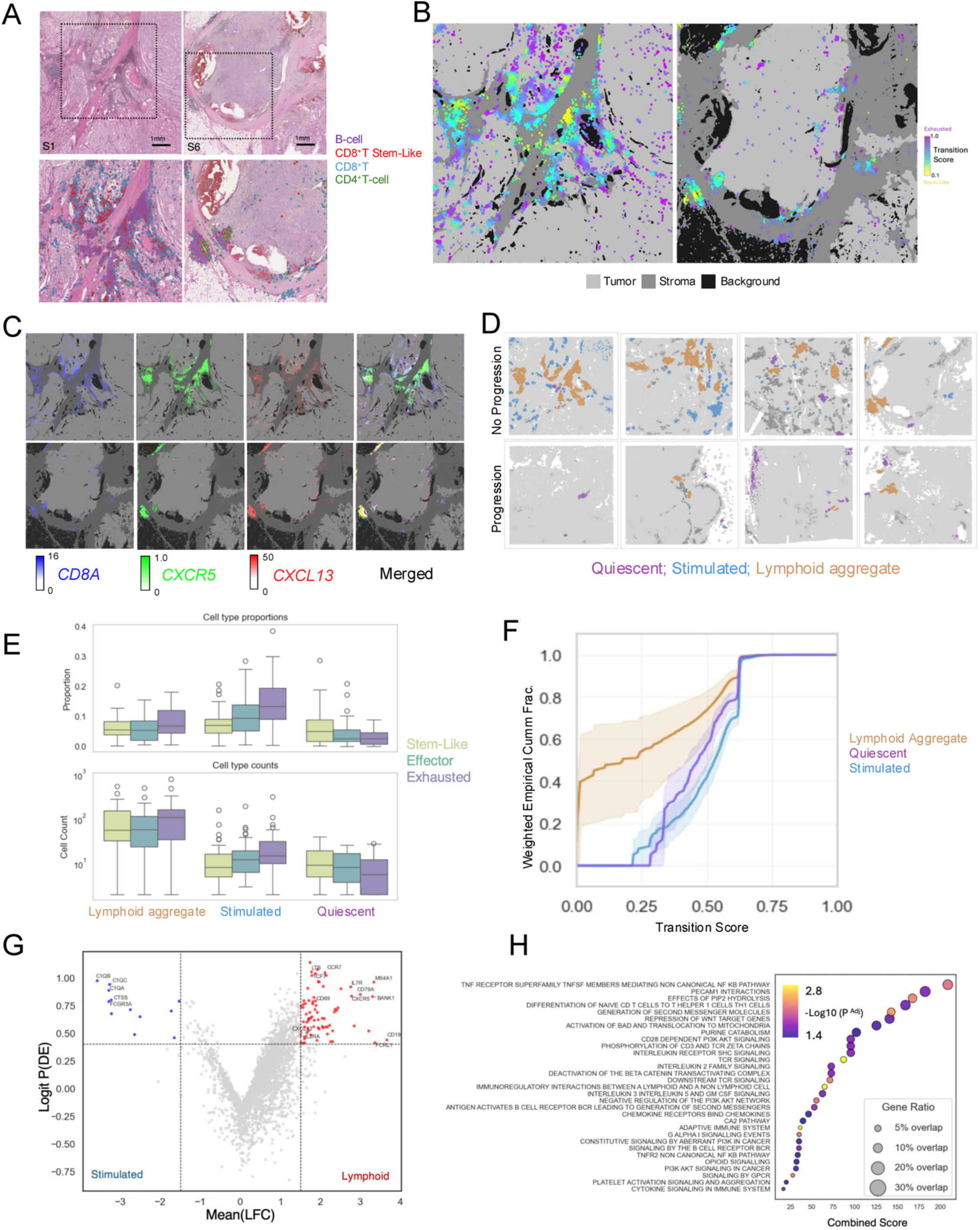
Spatial organization and functional characterization of stem-like CD8⁺ T cell containing niches in ccRCC. **(A)** Representative H&E sections with regions of interest defined (upper), with corresponding lymphocyte cell-type Voronoi estimated polygons overlaid on H&E (lower), highlighting spatial organization of immune cell subsets, with stem-like CD8^+^T cells geometrically expanded to visualize in B cell rich areas. **(B)** Stroma and tumor areas are colored by different shades of gray. Cells are represented as nuclei centroid dots scored along a progenitor-to-exhausted transcriptional axis using a continuous projection of gene expression signatures. **(C)** Spatial expression of *CD8A, CXCR5,* and *CXCL13* shows co-localization within lymphocyte-enriched regions, consistent with organized immune microenvironments supporting CD8⁺ T cell positioning. **(D)** Immune aggregates segregate into lymphoid, quiescent T cell, and stimulated T cell predominant immune clusters, representing a continuum of immune activation within the tumor microenvironment. **(E)** Spatial clustering identifies distinct immune niches with divergent CD8^+^ T cell state compositions. Proportions and absolute counts of CD8^+^ T cell subtypes are shown across spatial aggregate groups, where individual points are immune aggregates. **(F)** CD8⁺ stem-like T cell scores are differentially distributed across spatial clusters, with enrichment in lymphoid aggregates. ECDFs highlight shifts in stem-like populations, with 5–95% confidence intervals derived from bootstrap resampling. Scores are additionally weighted for visualization to emphasize low-abundance stem-like states; this weighting does not affect statistical comparisons between clusters. **(G)** Volcano plot showing comparative gene expression analysis of T cell–enriched clusters identifies distinct transcriptional programs associated with lymphoid-like niches. Subset of genes with high probability of differential expression, including *CXCR5* and *TCF7*, are annotated on the volcano plot. **(H)** Gene set enrichment analysis (Reactome C2) reveals pathway-level differences across spatial clusters, highlighting reactome terms associated with lymphoid aggregates compared to other immune cells. Dot size reflects gene set overlap, and color indicates −log10 false discovery rate.

To formalize these observations, we classified immune aggregates based on cellular composition and *CXCR5/CXCL13*-associated features. We identified three major aggregate classes, each containing at least 20% T cells, and mapped these across tumor sections based on CD8^+^ T cell states and *CXCL13/CXCR5* expression (**Fig. 6D**). Aggregates termed “Quiescent” lacked *CXCL13-CXCR5* co-expression and were characterized by CD8⁺ T cells with low exhaustion. “Stimulated” aggregates also lacked *CXCL13-CXCR5* co-expression but contained a higher proportion of exhausted CD8⁺ T cells. In contrast, “Lymphoid” aggregates showed *CXCL13-CXCR5* co-expression and were enriched for stem-like CD8⁺ T cells and B cells (**Fig. 6E; Fig. S13**). Cumulative stem-to-exhaustion transition scores further showed that Lymphoid aggregates contained a greater fraction of CD8⁺ T cells in lower transition states, consistent with enrichment for less differentiated, i.e. stem-like, T cell phenotypes (**Fig. 6F**).

Across slides, immune aggregates in progression samples were smaller and contained fewer cells than those in non-progression samples (median difference [non-progression − progression] = 70.14; p = 0.029). Because aggregates were relatively sparse and small in progression samples, downstream comparative analyses focused on dominant aggregate classes in non-progression tumors. Comparison of gene expression between lymphoid aggregates and other immune cell populations identified enrichment of *LTB*, markers of undifferentiated B cells, and several genes associated with stem-like T cell states (*TCF7, IL7R, CCR7, IL2RA*) in Lymphoid aggregates (**Fig. 6G**). Over-representation analysis similarly showed enrichment for pathways related to T cell co-stimulation, cytokine signaling, differentiation, and antigen presentation (**Fig. 6H; Table S8**). Collectively, these data support a model in which CXCL13-CXCR5–associated Lymphoid aggregates represent more activated, immunologically interactive niches with features consistent with tertiary lymphoid structure–like organization.

### CXCL13 suppresses tumor growth, enriches stem-like CD8⁺ T cells, and associates with improved clinical outcomes in ccRCC

Given the spatial enrichment of *CXCR5* expressing stem-like CD8⁺ T cells within CXCL13-associated lymphoid niches, we next assessed whether tumor-derived CXCL13 functionally shapes the intratumoral CD8⁺ T cell compartment and influences tumor growth *in vivo*. We used a syngeneic RENCA tumor model with wild-type (WT), CXCL13-overexpressing (CXCL13 High), and CXCL13 knockout (CXCL13 KO) cell lines. CXCL13 overexpression suppressed tumor growth with minimal tumor expansion and reduced tumor mass compared with WT and CXCL13 KO tumors (**Fig. 7A**). Conversely, CXCL13 KO tumors demonstrated accelerated growth, suggesting tumor-derived CXCL13 modulates tumor progression.

**Figure 7.**
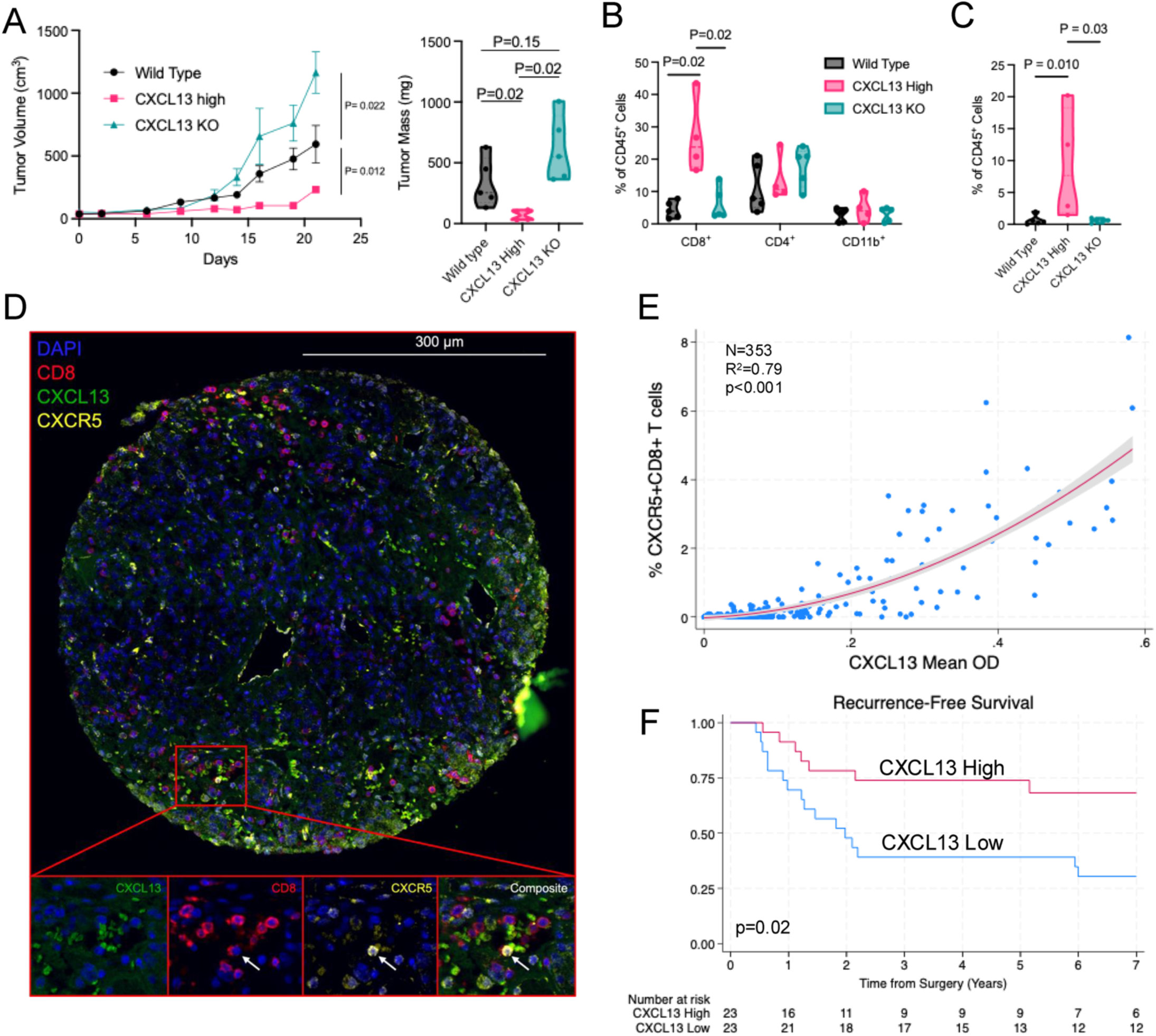
CXCL13 overexpression suppresses RENCA tumor growth, enriches progenitor-like CD8⁺ T cells in vivo, and associates with CXCR5⁺ CD8⁺ T cells and clinical outcomes in human high risk non-metastatic ccRCC. **(A)** Tumor growth kinetics (left) and final tumor mass (right) in BALB/c mice following subcutaneous injection of RENCA wild-type (WT), CXCL13-overexpressing (CXCL13 High), or CXCL13 knockout (CXCL13 KO) cells. Tumor volume was measured every 3 days for 3 weeks. (N = 5 mice per group). **(B)** Flow cytometric quantification of intratumoral immune populations, shown as percentage of CD45⁺ cells, including CD8⁺ T cells, CD4⁺ T cells, and CD11b⁺ myeloid cells. N=4-5 tumors per group. One tumor in the CXCL13 High cohort was insufficient quality for flow cytometry. **(C)** Frequency of CD8⁺CXCR5⁺TCF1⁺ cells within the CD45⁺ compartment across tumor groups. N=4-5 tumors per group. For panels A–C, p values were calculated using two-sided Mann–Whitney tests. **(D)** Representative multiplex immunofluorescence images of ccRCC TMA cores stained for CD8 (magenta), CXCR5 (cyan), and CXCL13 (green). Arrow demonstrates a CD8+CXCR5+ T cell. **(E)** Correlation between CXCL13 expression (mean optical density) and the proportion of CXCR5⁺CD8⁺ T cells across individual tumor cores (n = 353 cores from 46 tumors). Data are shown with a quadratic fit and 95% confidence band. A quadratic model provided a superior fit compared with a linear model (R² = 0.79 vs 0.76; likelihood-ratio test p < 0.001), with significant linear and quadratic terms (both p < 0.001). **(F)** Recurrence-free survival of patients with non-metastatic ccRCC treated with surgery stratified by tumor CXCL13 expression (high vs low), defined by the median CXCL13 optical density across sampled regions per tumor. N = 23 “CXCL13 High”, N = 23 “CXCL13 Low”. P value calculated using the log-rank test.

We next examined the impact of CXCL13 on the RENCA tumor immune microenvironment. CXCL13 overexpression was associated with increased intratumoral CD8⁺ T cells, whereas differences in CD4⁺ T cells and CD11b⁺ myeloid populations were less pronounced (**Fig. 7B**). Notably, CXCL13 High tumors exhibited a higher proportion of CD3⁺CD8⁺CXCR5⁺TCF1⁺ T cells within the CD45⁺ compartment than WT and CXCL13 KO tumors, consistent with enrichment of stem-like CD8⁺ T cells (**Fig. 7C; Fig. S14**).

To evaluate this relationship in human ccRCC, we analyzed a TMA cohort of 46 patients with high-risk, non-metastatic disease (**Table 1**). Multiplex immunofluorescence demonstrated a correlation between CXCL13 expression and CXCR5⁺CD8⁺ T cells within tumor regions (R² = 0.79, p<0.001), supporting an association between CXCL13 and CXCR5⁺CD8⁺ T cell infiltration in human tumors (**Fig. 7D,E**). Finally, to assess clinical relevance, we examined recurrence-free survival according to tumor CXCL13 expression. Patients with CXCL13-high tumors had improved recurrence-free survival compared with those with CXCL13-low tumors (log-rank p=0.02), supporting CXCL13 as a prognostic biomarker in high-risk ccRCC (**Fig. 7F**).

**Table 1.** Clinicopathologic characteristics of the human tissue microarray (TMA)

| **TMA Cohort** | **No Recurrence**<br>**(N=20)** | **Recurrence**<br>**(N=26)** | **P value** |
| --- | --- | --- | --- |
| **Median age, years (IQR)** | 61 (51-70) | 57 (50-66) | 0.3 |
| **Female sex, no. (%)** | 8 (40) | 9 (35) | 0.8 |
| **Pathologic T-stage, no.**<br>**(%)** |  |  | 0.2 |
| T2 | 10 (50) | 8 (31) |  |
| T3-T4 | 10 (50) | 18 (69) |  |
| **Median pathologic tumor**<br>**diameter, cm (IQR)** | 9 (7.2-9.4) | 9.1 (8-13) | 0.2 |
| **Grade, no. (%)** |  |  | 0.4 |
| 1-2 | 11 (55) | 10 (38) |  |
| 3-4 | 9 (45) | 16 (62) |  |
| **Median Follow-up, years**<br>**(IQR)** | 11 (8-15) | 7 (5-11) | 0.08 |

## DISCUSSION

In this study, we identify CXCL13 as a functional regulator of CD8^+^ T cell recruitment and spatial immune organization in ccRCC. Across human tumor profiling, migration assays, single-cell and spatial transcriptomic analyses, and *in vivo* models, our data provide evidence that CXCL13 helps recruit CXCR5^+^CD8^+^ T cells within lymphoid-like stromal niches enriched for less differentiated, stem-like states. These niches were characterized by coordinated CXCL13-CXCR5 expression, B cell enrichment, and transcriptional programs linked to costimulation, cytokine signaling, and antigen presentation. Tumor-derived CXCL13 also suppressed murine tumor growth, increased intratumoral CD8^+^ T cells, enriched CXCR5^+^TCF1^+^CD8^+^ T cell populations, and was associated with improved recurrence-free survival in patients with high-risk non-metastatic ccRCC.

These findings build on prior studies establishing that TCF1-associated stem-like, or progenitor, CD8^+^ T cells may serve as an intratumoral reservoir that sustains downstream effector and terminally exhausted populations [12–14, 17]. Our data suggest that CXCL13 plays a key role in how these cells are recruited to and spatially organize within ccRCC tumors. In multiple migration systems, CXCL13 promoted CD8^+^ T cell trafficking, enriched CXCR5^+^ proportions among migrating CD8^+^ T cells, and reduced migration after blockade of either CXCL13 or CXCR5. Together, these findings support the interpretation that CXCL13 is not merely a marker of an inflamed microenvironment, but an active component of CD8^+^ T cell recruitment in ccRCC [15, 16, 33].

Our single-cell and spatial analyses further suggest that CXCR5 and CXCL13 mark related but distinct positions along a continuous CD8^+^ T cell differentiation landscape. CXCR5 associated more strongly with stem-like and resident memory programs, whereas CXCL13 aligned with later differentiation states including cytotoxicity and exhaustion pathways. These findings support a model in which CXCL13-rich niches recruit or retain CXCR5-associated stem-like CD8^+^ T cells, some of which may then differentiate locally into more activated or exhausted states within the tumor [11, 30, 33]. This interpretation is supported by spatial data, in which stem-like CD8^+^ T cells were enriched in structured immune aggregates with B cells and coordinated CXCL13-CXCR5 expression rather than being diffusely distributed throughout the tumor. More broadly, this framework fits with prior niche-based models in which less differentiated stem-like CD8^+^ T cells enter tumors and acquire fuller effector features locally [7, 11, 34].

An additional insight from our study is that CXCL13 may mark a favorable immune ecosystem even though it is also associated with more differentiated or exhausted CD8^+^ T cell states. Our data suggest that this is not necessarily contradictory. Rather, CXCL13 may function at two levels which is supported by prior studies: as a chemokine that helps recruit CXCR5-associated stem-like CD8^+^ T cells [33], and as a feature of more differentiated CD8^+^ states [30]. Together, these observations suggest that CXCL13-rich niches may help recruit or retain a stem-like CD8^+^ compartment while also reflecting local differentiation into downstream activated or exhausted states.

Our findings should be interpreted in the context of several limitations. CXCR5 expression alone does not define a single CD8^+^ T cell identity, and CXCR5^+^CD8^+^ T cells are observed to have biologically heterogeneous functions across disease settings [35]. Accordingly, our interpretation relies less on CXCR5 as a standalone feature of a distinct CD8+T-cell state and more on its concordance with TCF7 and IL7R expression along inferred pseudotime dynamics, lower dimensional representations of holistic gene expression, and spatial localization patterns. In addition, the human cohorts are modest in size, particularly for spatial profiling, and our observational analyses cannot fully establish lineage relationships or causality in human tumors. However, the convergence of migration assays, human tissue data, single-cell and spatial analyses, and the RENCA murine model supports a consistent biologic framework.

Overall, our findings position the CXCL13-CXCR5 axis as a mechanism linking immune recruitment, niche organization, and a favorable stem-like CD8^+^ T cell compartment in ccRCC. More broadly, they suggest that effective anti-tumor immunity may depend not only on the presence of intratumoral CD8^+^ T cells, but on the local recruitment and maintenance of the specific CD8^+^ states capable of sustaining that response.

## Acknowledgements

The authors would like to thank the University of Wisconsin Biotechnology Gene Expression Center for assistance with spatial transcriptomics. The authors also wish to thank the University of Wisconsin Translational Research Initiatives in Pathology (TRIP) Laboratory, supported by the UW Department of Pathology and Laboratory Medicine, UWCCC (P30 CA014520) and the Office of The Director-NIH (S10 OD023526) for the use of their facilities.

## Author Contributions

Conceptualization: D.D.S., K.D.N., E.J.A.; Data Curation: D.D.S., K.D.N., M.L.; Formal Analysis: D.D.S., K.D.N., M.L.; Funding acquisition: D.D.S.; Methodology: D.D.S., K.D.N., M.L., K.E., S.C.K, E.J.A.; Project Administration: D.D.S., E.J.A.; Resources: D.D.S., K.D.N., E.N., S.M.M., S.C.K., C.M.C., E.J.A.; Software: K.D.N.; Supervision: D.D.S., E.J.A.; Visualization: D.D.S., K.D.N., M.L.; Writing original draft: D.D.S., K.D.N., E.J.A.; Writing-review and editing: D.D.S., K.D.N., M.L., P.M., Y.L., Y.Z., R.H., W.H., K.E., T.K., P.L., D.F.R., E.N., S.M.M., D.J.B., S.C.K., C.M.C., E.J.A.

## Data Availability

Single cell sequencing data from the Li et al. and Krishna et al. are publicly available. All data generated in this study are available upon request from the corresponding author.

## Ethics Approval

This study was performed in line with the principles of the Declaration of Helsinki. Approval for human subject work was granted by the Ethics Committee of the University of Wisconsin (#2024-0997). Approval for investigations involving mice was granted by the University of Wisconsin-Madison Institutional Animal Care and Use Committee (#M006825-A02).

## Consent to Participate

For all research involving human subjects, freely-given, informed consent to participate in this study was obtained from all participants. All participants were ≥18 years of age.

