## Supplementary Figures for "CXCL13-CXCR5 Signaling in CD8⁺ T Cell Recruitment and Lymphoid Immune Organization in Clear Cell Renal Cell Carcinoma"

| Supplementary Figures | Page Number |
| --- | --- |
| Figure S1 | 2 |
| Figure S2 | 3 |
| Figure S3 | 4 |
| Figure S4 | 5 |
| Figure S5 | 6 |
| Figure S6 | 7-8 |
| Figure S7 | 9 |
| Figure S8 | 10 |
| Figure S9 | 11 |
| Figure S10 | 12 |
| Figure S11 | 13-14 |
| Figure S12 | 15 |
| Figure S13 | 16 |
| Figure S14 | 17 |

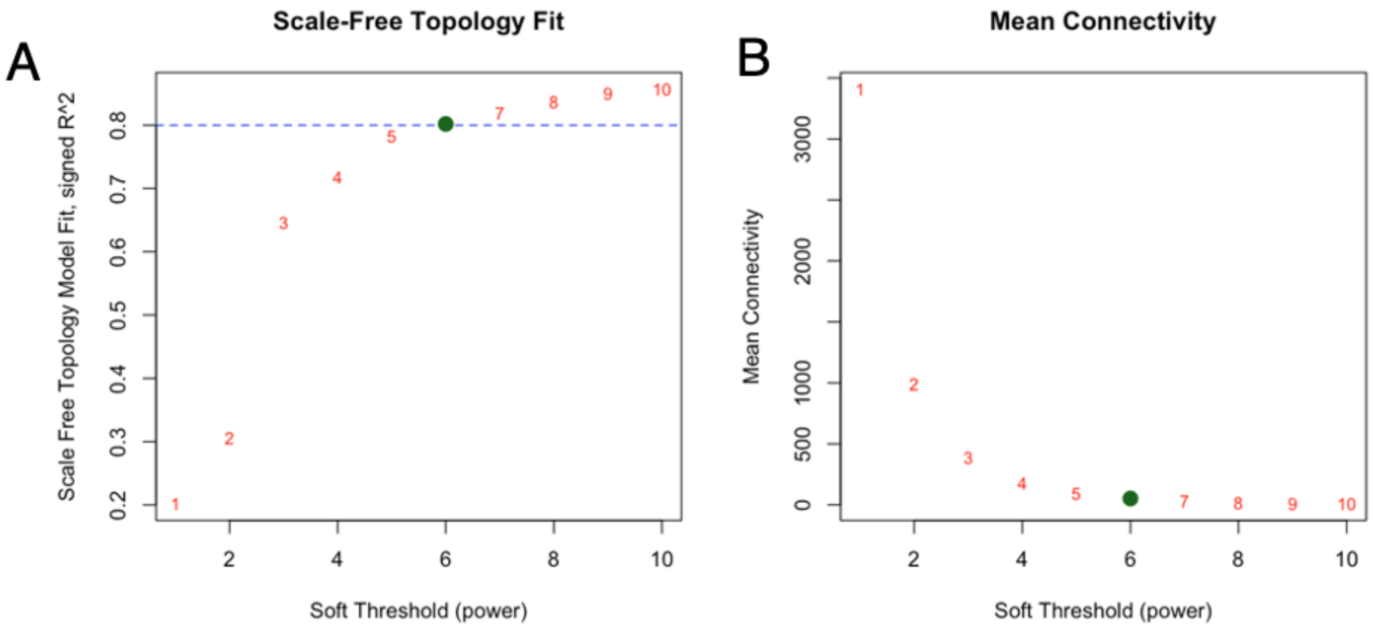


**Supplementary Figure S1. Selection of Weight Gene Co-Expression Network Analysis soft-thresholding power.(A)** Scale-free topology fit index (signed $R^2$) across candidate soft-thresholding powers ($\beta$). The dashed line indicates the heuristic threshold $R^2 = 0.8$, commonly used to select a network in which the connectivity distribution approximately approximates a power-law on a log–log scale, consistent with the scale-free topology assumption used in biologically interpretable gene co-expression networks. **(B)** Mean network connectivity (average node degree) as a function of $\beta$, reflecting changes in network sparsity with increasing thresholding power. The selected power ($\beta = 6$) represents the smallest value that satisfies the scale-free topology criterion ($R^2 > 0.8$) while maintaining sufficient connectivity for robust and biologically meaningful co-expression structure.

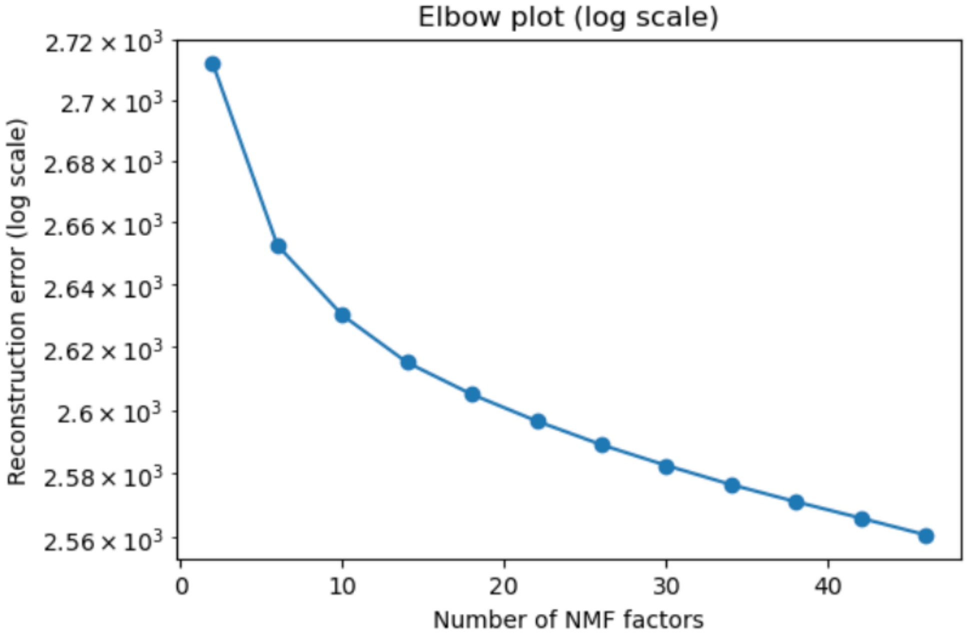


**Supplementary Figure S2. Selection of the optimal number of consensus non-negative matrix factorization (cNMF) components based on reconstruction error.** Reconstruction error (log scale) across a range of cNMF component counts. Increasing the number of components improves model fit, as reflected by decreasing reconstruction error; however, gains plateau beyond the elbow point, indicating an optimal balance between model complexity and explanatory power.

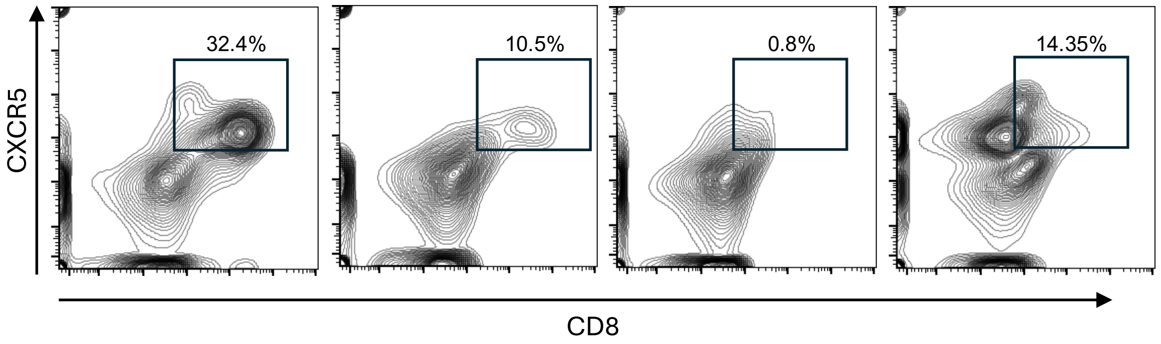


**Supplementary Figure S3. Representative flow cytometry plots of CXCR5 expression among tumor-infiltrating CD8⁺ T cells.** Representative contour plots demonstrating CXCR5 expression on CD8⁺ T cells isolated from individual ccRCC tumors corresponding to the quantification shown in Fig. 3A. Tumor tissues were dissociated into single-cell suspensions using enzymatic digestion (collagenase, DNase, and hyaluronidase) with a gentleMACS dissociator. CD8⁺ T cells were isolated from tumor tissue using positive selection with REAlease CD8 microbeads, and purity (>90%) was confirmed by flow cytometry. Cells were gated on live, singlet lymphocytes followed by CD8⁺ T cells, and CXCR5 expression is displayed on the y-axis versus CD8 on the x-axis. Boxes indicate the CXCR5⁺ CD8⁺ T cell population, with percentages shown for each representative sample.

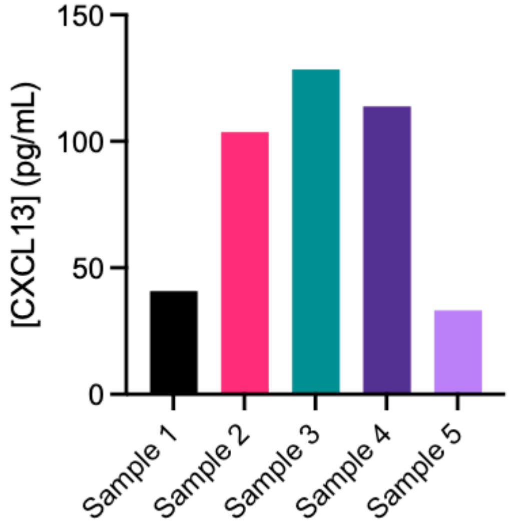


**Supplementary Figure S4. CXCL13 secretion by primary tumor-derived ccRCC cell cultures**. CXCL13 concentrations from five independent primary ccRCC cell cultures were quantified by ELISA demonstrating expression within all cell cultures.

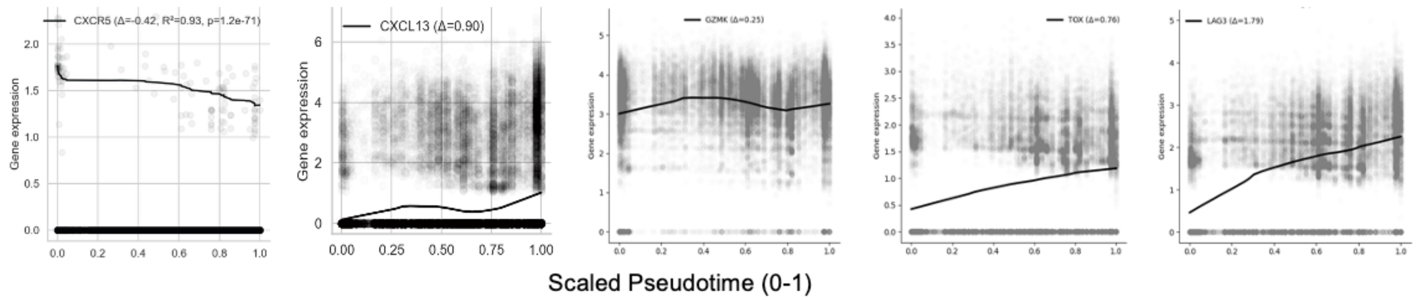


**Supplementary Figure S5. Pseudotime dynamics of selected genes in ccRCC.** Gene expression trends along cells ordered by pseudotime, including *CXCR5, CXCL13, GZMK, TOX,* and *LAG3*. Curves represent LOWESS-smoothed fits of library size normalized expression across scaled min-max scaled pseudotime. For the *CXCR5* plot, LOWESS was estimated from non-zero values, while still showing all *CXCR5* expression values for transparency.

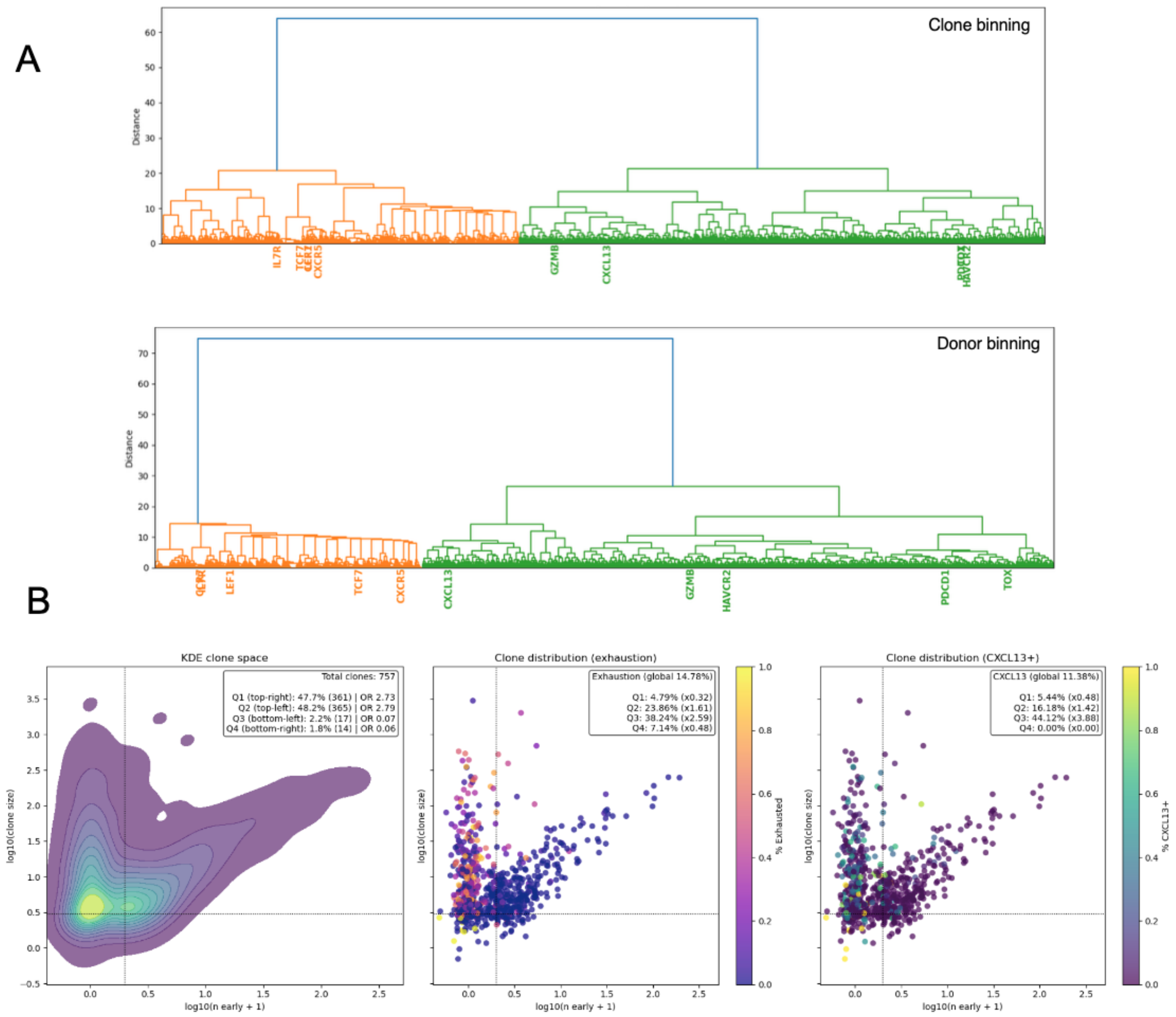


**Supplementary Figure S6. Clonotype-resolved analysis of stem-like gene expression patterns and clonal composition in CD8⁺ T cells (A)** Dendrograms showing gene expression patterns across binned Pseudotime to prevent overdominance of specific clones or donors. Genes cluster into two modules with distinct trajectories and are displayed as hierarchical trees. Top: cells grouped by pseudotime bin and clone identity. Bottom: cells grouped by pseudotime bin and donor. Highlighted genes mark key clusters. Vertical lines indicate clones containing ≥1 stem-like cell; horizontal line marks clones with ≥3 total cells. **(B)** Left: relationship between clone size and number of stem-like CD8⁺ T cells per clone shown as kernel density plot. Middle: same relationship as scatter plot showing points colored by the fraction of exhausted $CD8^+$ T cells per clone. Right: same plot colored by percent $CXCL13^+$ cells. Insets show the distribution of exhausted-cell fractions and clone densities across quadrants.

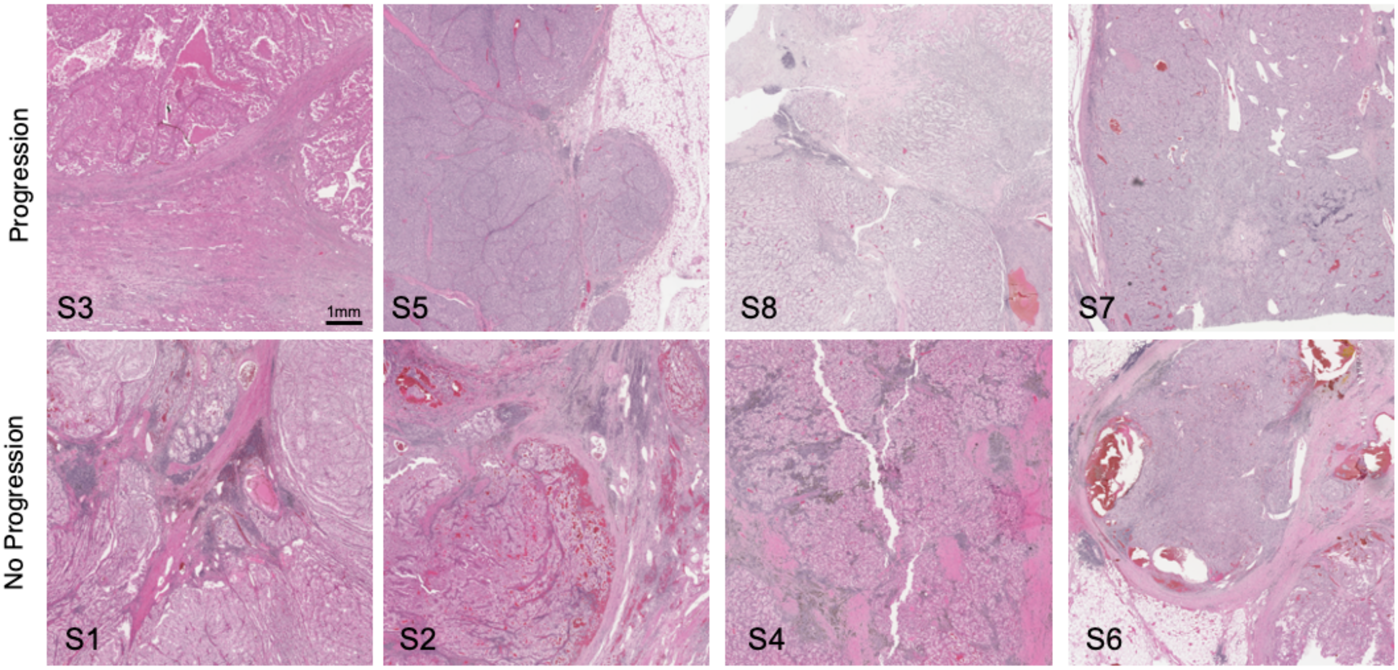


**Supplementary Figure S7. Representative H&E sections corresponding to spatial transcriptomic capture areas.** Representative hematoxylin and eosin (H&E)–stained sections from ccRCC tumors demonstrating the regions selected for spatial transcriptomic profiling. Images are aligned to the sequenced capture areas shown in Fig 5A. Samples are shown across progression and no progression groups.

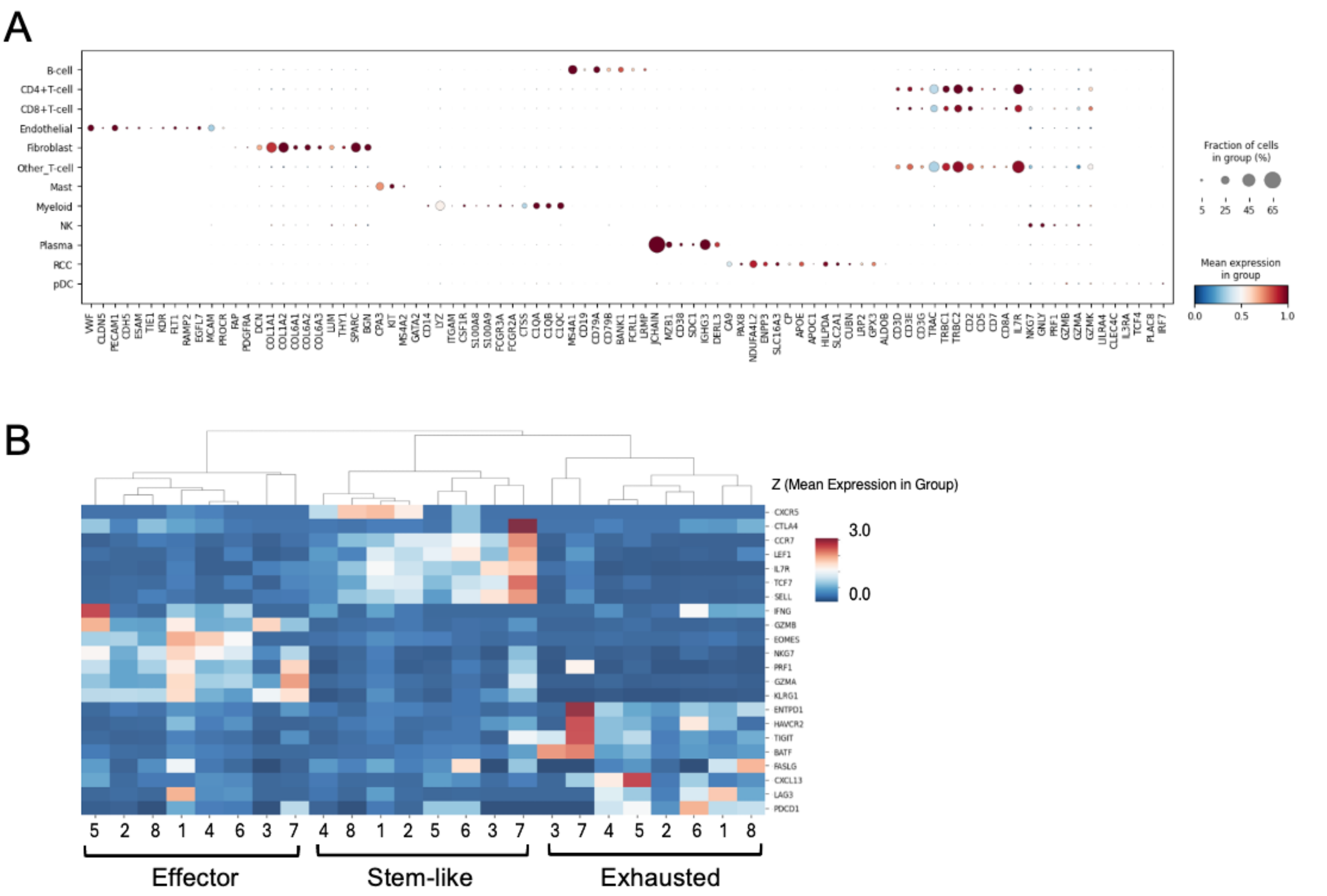


**Supplementary Figure S8. Validation of spatial cell-type and CD8$^+$ T cell state annotations. (A)** Dot plot showing expression of canonical marker genes across annotated cell types derived from spatial transcriptomic data, confirming expected lineage-specific expression patterns. Dot size represents the percentage of cells expressing each gene, and color indicates mean expression, scaled per gene. **(B)** Heatmap showing the average expression of selected CD8$^+$ T-cell marker genes derived from scRNA-seq data. Gene ordering is based on hierarchical clustering of Z-scored mean expression values, following variance stabilization of single cell estimated expression values. Samples are stratified into effector, stem-like, and exhausted CD8$^+$ T cell groups. These three functional states form distinct clusters based on their characteristic marker gene expression profiles, supporting the criteria used for CD8$^+$ T cell annotation.

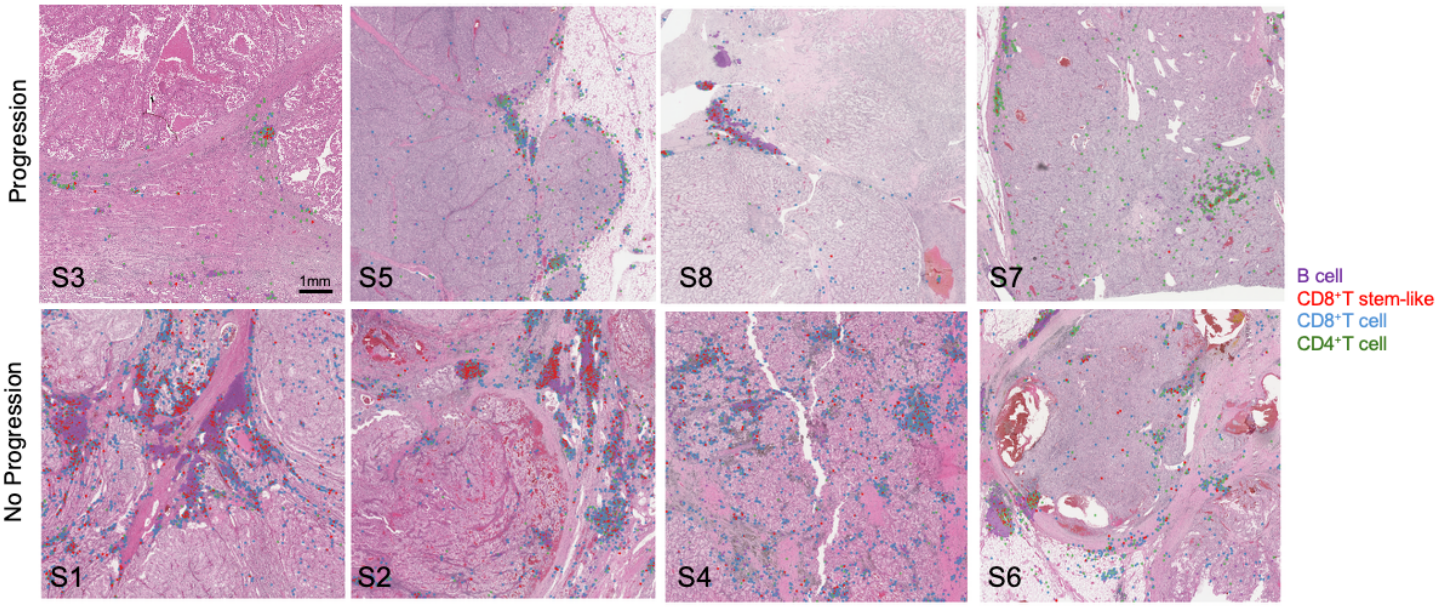


**Supplementary Figure S9. Spatial distribution of lymphocyte subsets across all analyzed ccRCC tumor sections.** Representative hematoxylin and eosin (H&E)–stained sections from all analyzed tumors, with spatial transcriptomic cell annotations overlaid. Cells are colored by broad lymphocyte subsets, including B cells, $CD4^{+}$T cells, $CD8^{+}$ T cells, and *CXCR5*-expressing stem-like $CD8^{+}$ T cells, as indicated in the legend. Samples are grouped by progression status (progression vs no progression). ENACT-style polygons labeled as $CD8^{+}$T cell stem-like were expanded for visualization purposes. These images illustrate the heterogeneous spatial distribution of lymphocyte populations across tumor sections and provide context for the representative regions shown in Fig 6A.

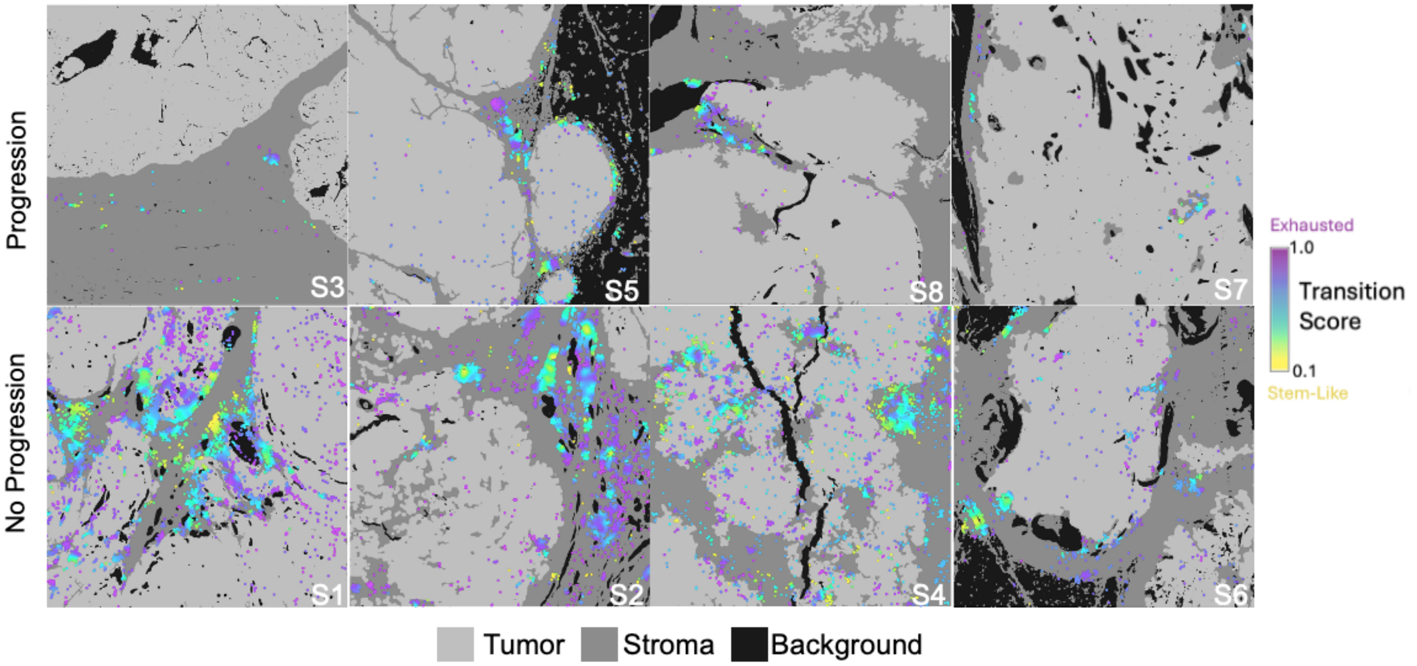


**Supplementary Figure S10. Spatial distribution of CD8$^{+}$ T cell stem–exhaustion scores across ccRCC tumor sections.** Gaussian-smoothed spatial maps of CD8$^{+}$ T cell stem-like/exhaustion scores overlaid on tumor sections from all analyzed samples. These maps reveal heterogeneous spatial patterns of inferred CD8$^{+}$ T cell states, ranging from stem-like to more exhausted phenotypes, across ccRCC tumors. This cohort-wide view provides context for the representative regions shown in Fig. 6C. Tumor sections are grouped by tumors that progressed or did not progress to metastatic disease.

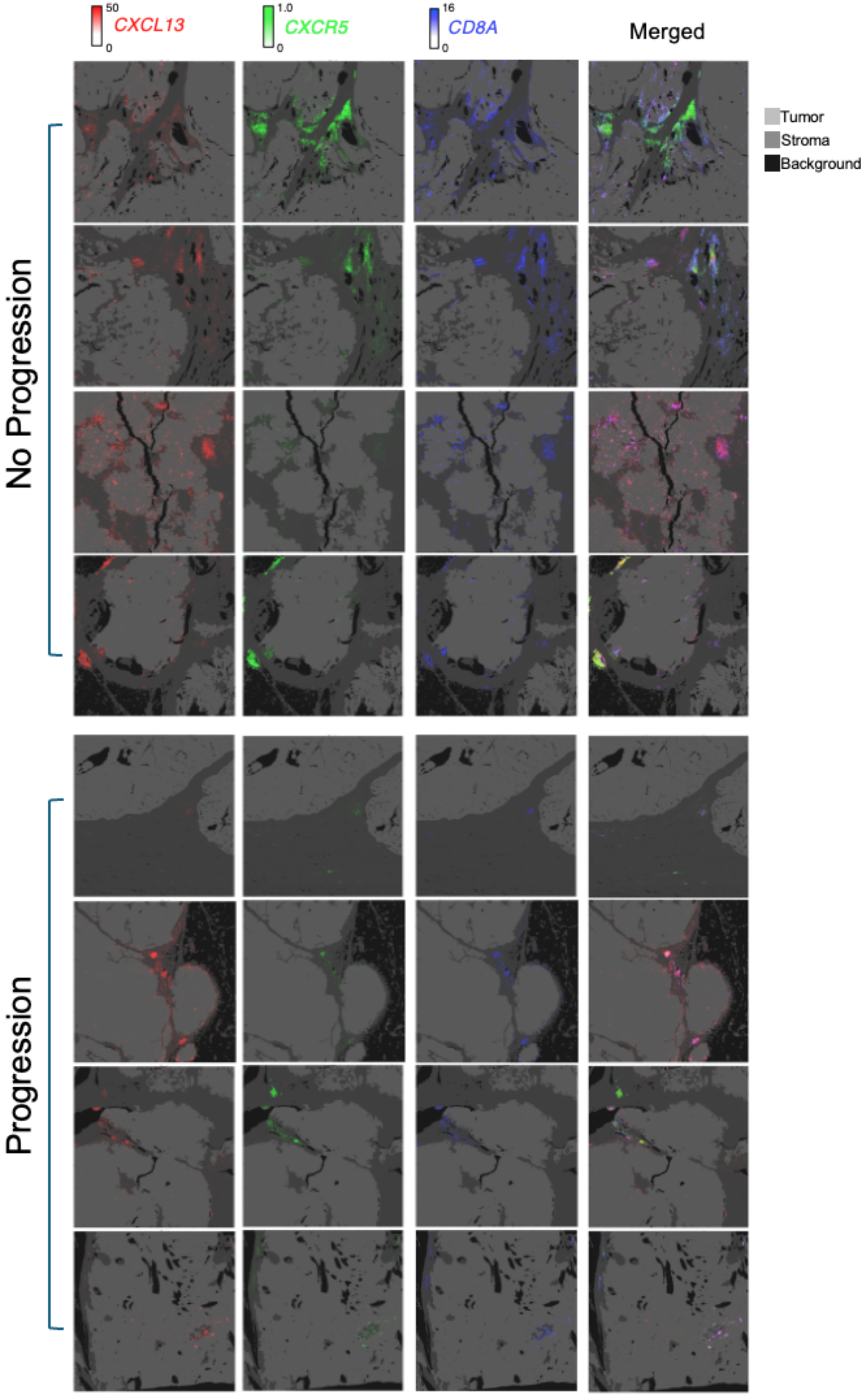
50
CXCL13
0
1.0
CXCR5
0
16
CD8A
0
Merged
Tumor
Stroma
Background
No Progression
Progression

**Supplementary Figure S11. Spatial distribution of *CXCL13, CXCR5,* and *CD8A* estimated expression across ccRCC tumor sections.** Spatial transcriptomic maps of *CXCL13, CXCR5,* and *CD8A* scVI-normalized expression overlaid on tumor sections across all analyzed samples. Gene expression values were variance-stabilized and clipped to the 1st–99th percentile range for visualization. These maps demonstrate heterogeneous spatial expression patterns of *CXCL13* and *CXCR5* and highlight regions of co-localization with $CD8^+$ T cells. This figure provides cohort-wide context for the representative regions shown in Fig. 6B. Tumor sections are grouped by tumors that progressed or did not progress to metastatic disease.

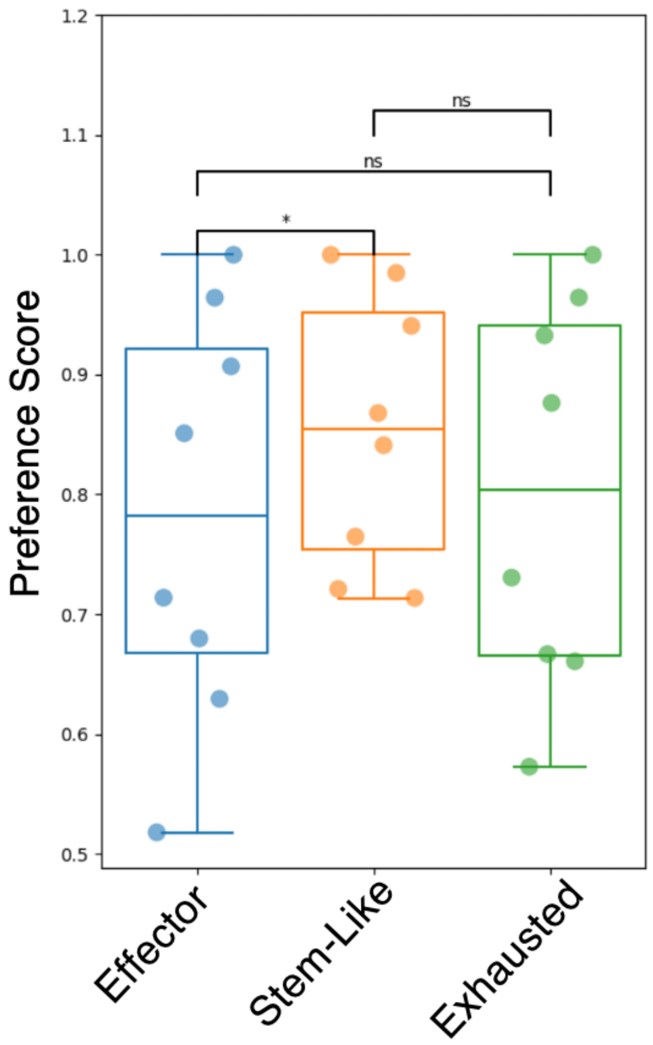


**Supplementary Figure S12. Pairwise comparison of spatial preference scores across CD8⁺ T cell groups.** Each point represents the slide-level difference in stromal preference for each of the three CD8⁺ T cell states, computed as the fraction of spots assigned to stroma relative to total spots in stroma and tumor compartments. Boxplots summarize distributions across slides. Statistical significance was assessed using a paired Wilcoxon signed-rank test (* $p < 0.05$, ns=not significant).

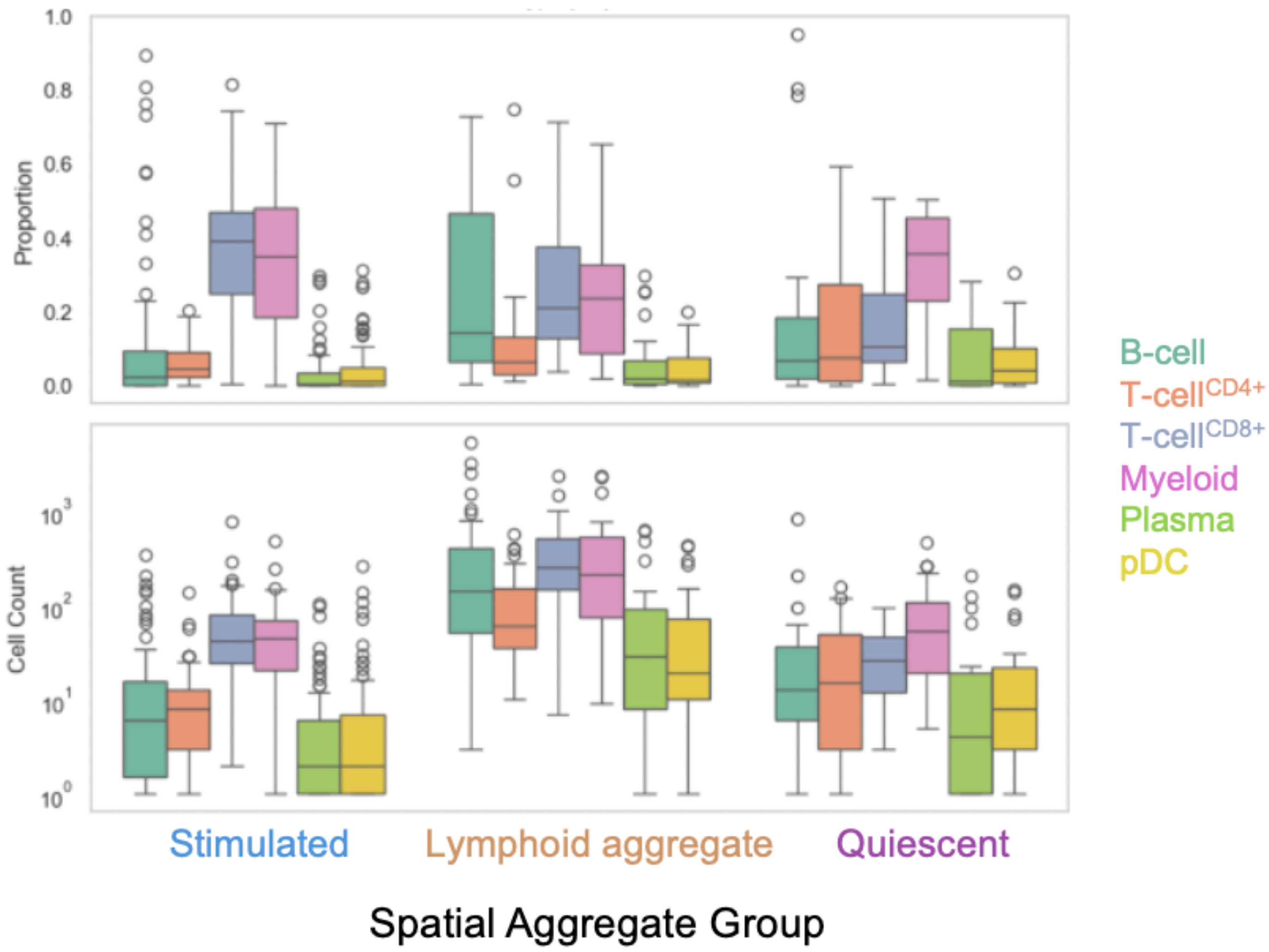


**Supplementary Figure S13. Spatial clustering identifies distinct immune niches with divergent cellular composition.** Proportions and absolute counts of major immune populations are shown across spatial aggregate groups (stimulated, lymphoid aggregate, and quiescent), with rare cell types (such as MAIT, NK, gamma T-cells) excluded.

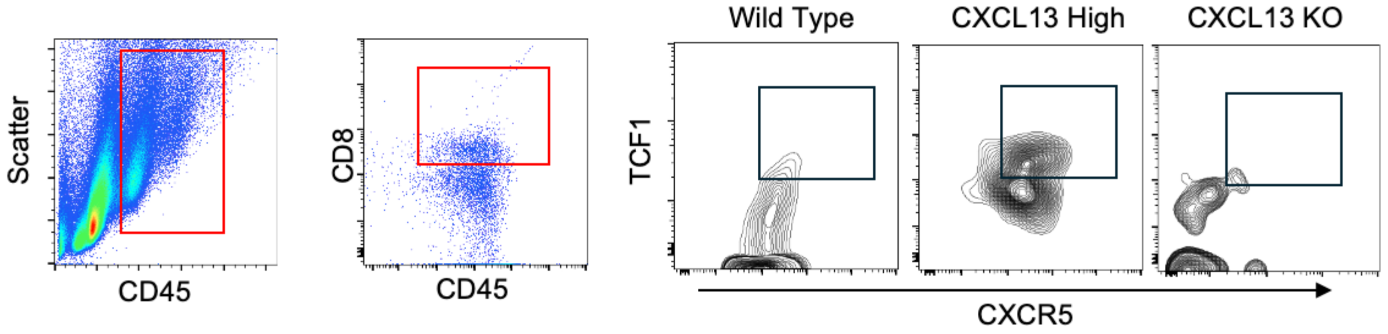


**Supplementary Figure S14. Flow cytometry gating strategy for identification of CXCR5⁺TCF1⁺ CD8⁺ T cells in RENCA tumor models.** Representative flow cytometry plots demonstrating the gating strategy used for analysis of tumor-infiltrating immune populations in RENCA tumors expressing wild type, CXCL13-overexpressing (CXCL13 High), or CXCL13 knockout (CXCL13 KO) cells. Single-cell suspensions were generated from tumors, and live leukocytes were first identified by gating on $CD45^+$ cells. From the $CD45^+$ population, $CD8^+$ T cells were selected based on CD8 expression. Within $CD8^+$ T cells, CXCR5 and TCF1 expression were assessed to define the $CXCR5^+TCF1^+$ subset. Boxes indicate gated populations at each step. These gating strategies correspond to the quantification of $CD45^+CD8^+CXCR5^+TCF1^+$ cells shown in Fig. 7 (main figure panels B and C).
