## Supplementary Methods for "CXCL13-CXCR5 Signaling in CD8⁺ T Cell Recruitment and Lymphoid Immune Organization in Clear Cell Renal Cell Carcinoma"

**SUPPLEMENTARY METHODOLOGY**

**RNA preparation and qPCR**

Total RNAs were prepared by Trizol reagent (Invitrogen). After reverse transcription, the resulting cDNA was used for quantitative PCR experiments (Bio-Rad) with SYBR green and the primers for variable cytokines. Sequences of used primers are below. The experiments were repeated three times, duplicates and independently (mean ±SEM).

hCXCL9: for; CTGTTCCTGCATCAGCACCAAC, rev; TGAACTCCATTCTTCAGTGTAGCA

hCXCL10: for; GGTGAGAAGAGATGTCTGAATCC. rev; GTCCATCCTTGGAAGCACTGCA

hCXCL11: for; AAGGACAACGATGCCTAAATCCC, rev; CAGATGCCCTTTTCCAGGACTTC

hCXCL12: for; CTCAACACTCCAAACTGTGCCC, rev; CTCCAGGTACTCCTGAATCCAC

hCXCL13: for; TATCCCTAGACGCTTCATTGATCG, rev; CCATTCAGCTTGAGGGTCCACA

hCCL2: for; AGAATCACCAGCAGCAAGTGTCC, rev; TCCTGAACCCACTTCTGCTTGG

hCXCL13: for; TATCCCTAGACGCTTCATTGATCG, rev; CCATTCAGCTTGAGGGTCCACA

mCXCL13: for; CATAGATCGGATTCAAGTTACGCC, rev; GTAACCATTTGGCACGAGGATTC

**Tissue microarray**

A tissue microarray (TMA) was constructed using non-metastatic ccRCC tumors at high risk of recurrence (tumors ≥7cm in size at the time of surgery). Two cohorts of patients were included, those that did (N=26) or did not progress (N=20) to metastatic disease. These two patient cohorts were matched on age at surgery, gender, tumor size, tumor grade, and tumor stage. The TMA was constructed using 7-8 tumor cores per patient and built with a manual tissue arrayer (Beecher Instruments, Sun Prairie, WI, USA) with 0.6mm cores. The TMA was stained using multiplexed immunohistochemistry as previously described [1, 2]. 4μm thick TMA slides were deparaffinized and heat-induced epitope retrieval was carried out. Slides were stained for CXCL13 (ab246518), CD8 (Ventana #790-4460), CXCR5 (ab254415), and DAPI using a protocol previously described [1]. Slides were scanned using the Vectra 2 slide scanner (Akoya/Quanterix, MA) and an automated scanning protocol was created to acquire multispectral image cubes using the 20x objective. Control slides stained with only 1 chromogen were used to create a spectral library algorithm in Nuance v3.0.2 software (Akoya/Quanterix, MA). InForm v2.4 software was used to perform image analysis (Akoya/Quanterix, MA). The spectral library algorithm and image cubes were chosen to set up a workflow of algorithms of differentiation for tissue segmentation, cell segmentation and to measure biomarker expression. The algorithm workflow was applied to the full set of TMA image cubes. CXCL13 concentration was quantified as optical density. CD8^+^CXCR5^+^ T cells were quantified as the percentage of total nucleated cells in the tissue core. The correlation between CXCL13 and CD8^+^CXCR5^+^ T cells was evaluated using quadratic fit regression model. The association with tumor CXCL13 and recurrence-free survival probability was estimated for patients using the Kaplan-Meier method with follow-up measured from the time of surgery. The median CXCL13 density was measured across individual tumor cores per tumor and the median was calculated. Patients were stratified into either CXCL13 high or low cohorts based on the median density of each tumor being either higher or lower than the total cohort’s CXCL13 median density. Recurrence-free survival was compared between CXCL13 high and low cohorts using the log-rank test.

**Network-Based Analysis of Bulk RNA-seq Data from TCGA-KIRC via WGCNA**

Bulk RNA-seq data (log2(TPM + 1)) from The Cancer Genome Atlas (TCGA) M0 clear cell renal cell carcinoma (ccRCC) cohort were used for co-expression network analysis. Only patient samples with complete clinical annotations were retained, yielding a final dataset of n = 369. TPM (transcripts per million) normalization accounts for sequencing depth and gene length, enabling comparison of relative gene expression levels across samples.

Weighted gene co-expression network analysis (WGCNA) was performed using the WGCNA R package. A soft-thresholding power (β) was selected using the pickSoftThreshold function (range 1–20) to approximate scale-free topology. The optimal power (β = 6) was defined as the lowest value achieving a scale-free topology fit index (R² > 0.8) while maintaining adequate mean connectivity (Fig. S1). A signed adjacency matrix was constructed from pairwise gene expression correlations and transformed into a topological overlap matrix (TOM) to enhance network robustness.

Genes were hierarchically clustered based on TOM dissimilarity using average linkage, generating a gene dendrogram. Modules of co-expressed genes were identified using dynamic tree cutting, with branches of the dendrogram adaptively pruned to define discrete modules. Closely related modules were further merged based on eigengene similarity using a defined cut height threshold, resulting in a set of refined co-expression modules.

**Gene Ontology Overrepresentation Analysis**

The grey60 WGCNA module, selected based on its inclusion of the chemokine genes CXCL13 and CXCL9, was subjected to enrichment analysis against Gene Ontology (GO) Biological Process gene sets from the Molecular Signatures Database (MSigDB C5). Overrepresentation analysis was performed using Fisher’s exact test, with multiple testing correction applied using the Benjamini–Hochberg method.

Significantly enriched GO terms were ranked by adjusted p-value (false discovery rate, FDR), and the top 20 terms were retained for visualization. Enrichment results were summarized using scaled enrichment scores, with dot size representing gene overlap and color indicating −log10(FDR), enabling interpretation of the biological processes associated with the grey60 co-expression module.

**Single-cell RNA-seq processing and integration**

Single-cell RNA-seq data from the publicly available datasets by Krishna et al. [3] and Li et al. [4] were processed using tools within the Scanpy ecosystem. Cells with >10% mitochondrial transcript content, fewer than 500 detected genes, or low-quality annotations (as defined by the original studies) were excluded. Genes expressed in fewer than 10 cells were removed.

Count matrices were library-size normalized to a fixed total (10,000 counts per cell) and log-transformed using a natural logarithm (log1p). Highly variable genes (HVGs) were identified using the Seurat v3 flavor implemented in Scanpy. This feature set was augmented by inclusion of predefined chemokine genes of interest (e.g., CXCR5, CXCL13) and additional marker genes identified via Wilcoxon rank-sum differential expression analysis performed on log-normalized data to capture features informative of published cell-state annotations.

Principal component analysis (PCA) was performed on the scaled HVG matrix. To correct for donor- and batch-specific effects across datasets, Harmony integration was applied to the PCA embeddings using both donor and batch as covariates, generating batch-corrected low-dimensional representations for downstream analyses. Uniform Manifold Approximation and Projection (UMAP) embeddings were then computed from the Harmony-corrected principal components for visualization.

Neighborhood graphs were constructed using the corrected embeddings, followed by clustering with the Leiden algorithm. Broad cell-type annotations were transferred from the Li et al. reference dataset using CellTypist. CD8⁺ T cells were further subset based on original author annotations. Within this compartment, transcriptional states (progenitor, effector, effector memory, and exhausted) were assigned by majority-vote labeling of Leiden clusters, guided by generalized annotations from Li et al.

**CXCR5⁺ CD8⁺ T cell Identification and Modeling**

CD8⁺ T cells were defined based on curated annotations from the original datasets. Identification of CXCR5-expressing cells is complicated by the fact that CXCR5 does not delineate a discrete cell type but instead exhibits graded and state-dependent expression across multiple CD8⁺ T cell subsets. As a result, binary classification of CXCR5⁺ cells are inherently sensitive to technical dropout and thresholding choices, and may not reflect a biologically distinct population.

To account for these properties, CXCR5 expression was modeled using complementary probabilistic approaches. A Gaussian mixture model (GMM) was applied to log-normalized expression values to partition cells into low- and high-expression components. In parallel, a zero-inflated negative binomial framework implemented in single cell variational inference (scVI) was used to model gene expression while accounting for dropout and overdispersion, yielding posterior estimates of CXCR5 expression probability for each cell. CXCR5⁺ cells were defined as those assigned to the expressing component in the GMM or exhibiting high posterior expression probability in the scVI model.

Given the continuous and state-associated nature of CXCR5 expression, these annotations were not interpreted as defining a discrete lineage, but rather as a pragmatic stratification to facilitate visualization and comparative analyses. Accordingly, CXCR5⁺CD8⁺ T cell labels were used for visualization purposes, including dot plots (Fig. 4B) and stacked bar plots summarizing cell-type proportions in spatial transcriptomics data (Fig. 5B).

**Co-expression Structure Inference via Consensus Non-negative Matrix Factorization**

We then aim to measure how closely other genes align with the CXCL13 and CXCR5 program profiles in lower dimensional spaced defined by non-negative matrix factorization (cNMF) to determine whether they converge on similar transcriptional states and functions across CD8⁺ T cells. Gene expression programs were identified using cNMF, which decomposes expression of an expanded set of highly variable genes from CD8⁺ T cells into a set of recurrent, non-negative additive components. These components represent latent transcriptional programs capturing coordinated variation in gene expression across cells. We used cNMF specifically because it provides a low-dimensional representation of shared transcriptional structure, reducing noise inherent to sparse single-gene measurements and avoiding the instability of pairwise gene–gene correlations in high-dimensional single-cell data. cNMF also acknowledges that CD8+T-cell states are not discrete cell types, but rather a continuum of states.

Unlike gene–gene correlation approaches, which estimate relationships directly in the presence of substantial sparsity, dropout, and indirect regulatory effects, cNMF identifies modules of co-varying genes that are consistently expressed together across cells. This yields a more robust representation of biological signal by aggregating information across genes into interpretable programs, rather than relying on pairwise relationships that can be dominated by technical noise or confounded by differences in cell state composition.

Each cNMF component was interpreted as a transcriptional program reflecting continuous axes of variation such as memory-like, migratory, or effector-associated states, consistent with prior applications of this framework. Rather than re-defining the full biological interpretation of individual components, which has been previously established in the source publications, we focused on leveraging cNMF loadings as a representation of transcriptional structure to assess the multi-program expression of chemokines across CD8⁺ T cells.

These programs were treated as shared components of transcriptional structure. Accordingly, each gene was represented as a continuous loading profile across all inferred programs, reflecting its relative contribution to each transcriptional axis. This representation enables comparison of genes in a shared latent space defined by coordinated transcriptional programs, rather than relying on noisy pairwise similarity metrics.

Gene-level similarity in program space was quantified by comparing normalized cNMF loading vectors using cosine similarity. Loading profiles of CXCR5 and CXCL13 were used as reference vectors, and all genes were ranked according to their similarity to these profiles across the full set of programs. Prior to similarity computation, gene-level loading vectors were L2-normalized to ensure that comparisons reflected alignment in loading patterns across programs rather than differences in magnitude. Similarity scores were optionally z-scored across genes to enable comparison on a standardized scale.

Genes exhibiting high similarity to CXCR5 or CXCL13 loading profiles were interpreted as sharing coordinated representation across the same transcriptional programs, indicating similar positioning within the latent program space defined by cNMF. This framework enables comparison of gene behavior based on shared structure across approximately 4,000 genes, rather than assignment to discrete categories or reliance on direct gene–gene correlation structure.

To assess functional convergence, gene set enrichment analysis was performed using Enrichr (via gseapy). Genes ranked by similarity to CXCR5 and CXCL13 loading profiles were tested against predefined transcriptional signatures, including exhaustion, cytotoxicity, progenitor-like, and tissue residency programs. This analysis evaluated whether genes with similar cNMF-derived program structure also converge on shared annotated biological functions.

To support these findings, gene–gene associations were additionally evaluated using multiple complementary approaches, including Spearman and Kendall correlation and zero-inflated negative binomial modeling within CD8⁺ T cells. MAGIC imputation was applied to mitigate sparsity, and associations were assessed across a range of diffusion time parameters (Table S3).

**Pseudotime and T cell receptor analysis**

To assess CD8⁺ T-cell differentiation trajectories and potential dysfunctional states in ccRCC, we performed pseudotime trajectory analysis. Stem-like CD8⁺ T-cell clusters were designated as root cells for trajectory inference using Monocle3 on Harmony-corrected UMAP embeddings. The UMAP space was used to construct the underlying k-nearest-neighbor graph, as it preserves local transcriptional relationships in a reduced-dimensional space, enabling robust ordering of cells along a continuous differentiation axis in sparse single-cell RNA-seq data. Cells were then ordered along the inferred pseudotime trajectory.

To characterize dynamic changes in gene expression, cells were ordered along this inferred continuum and expression patterns of genes of interest were evaluated across pseudotime. These profiles were visualized and summarized using LOWESS smoothing and polynomial regression to reduce technical noise and highlight continuous transcriptional changes across CD8⁺ T-cell states along the trajectory.

To assess robustness to donor effects, clone size, and clonal composition, we generated dendrogram-based summaries of gene expression dynamics across inferred pseudotime-ordered cells. Clones were defined by identical paired αβ TCR sequences. Gene expression was aggregated by averaging log-normalized values within pseudotime bins, with binning strategies designed to minimize overrepresentation from any single clone or donor. In one configuration, cells were grouped by clone identity and divided into 40 pseudotime bins with a minimum of 80 cells per clone; in an alternative configuration, cells were binned jointly by pseudotime and donor identity. Genes were hierarchically clustered using Ward linkage based on similarity in their Expression profiles were computed across the pseudotime axis, and genes of interest were used to define two main dynamic modules corresponding to progenitor-like and exhausted-like transcriptional programs. We then compared the behavior of these modules between the root (progenitor-enriched) cluster and cells at the terminal end of the inferred pseudotime trajectory to determine whether root cells show evidence of expansion and what downstream cell states they most closely transition into. We further assessed whether these patterns were robust to donor effects as well as variation in clone size and clonal composition.

To characterize clonal dynamics along the trajectory, we quantified the relationship between clonal expansion and cell-state composition. A density plot summarized the association between variance-stabilized clone size and the number of progenitor-like CD8⁺ T cells per clone, while a scatter plot showed individual clonotypes colored by the proportion of exhausted CD8⁺ T cells. Insets summarize the distribution of terminal differentiation-associated states across clones and the density of clones across quadrant-defined regimes, illustrating how expanded progenitor-enriched clones might relate to downstream exhausted-like states.

Finally, we evaluated the distribution of CXCL13⁺ CD8⁺ T cells across pseudotime and within clonotypes to further define their relationship to the inferred differentiation trajectory and to identify which downstream transcriptional states might expand from the root population.

**Spatial transcriptomics preprocessing (Visium HD)**

8 representative tumor samples were selected for inclusion in the spatial transcriptomic analysis using the Visium HD platform (10xGenomics). All patients had high risk non-metastatic ccRCC and underwent surgery with curative intent. Four patients ultimately progressed to metastatic disease after surgery and four patients did not progress to metastatic disease. Patients from each cohort were balanced in terms of age, gender, tumor size, pathologic stage, grade, sarcomatoid and rhabdoid features. Tumor sections were selected by a trained genitourinary pathologist (Y. Z.) and included representative sections of the overall tumor and included both tumor and stromal containing areas.

To enable accurate co-registration of sequencing data with tissue morphology without relying solely on fiducial marker placement in the CytAssist image, full-resolution H&E images were aligned to CytAssist images using shared morphological features. Corresponding histological landmarks were identified in the CytAssist images and matched to features in the full-resolution H&E images using the Scale-Invariant Feature Transform (SIFT) algorithm. An affine transformation was then estimated from these matched feature pairs and applied to the full-resolution H&E image to achieve spatial alignment. Registration accuracy between the H&E and CytAssist images was evaluated by manual inspection across multiple regions of the tissue to confirm consistent local and global alignment.

The aligned H&E and CytAssist images were then mapped to the Visium HD reference slide coordinate system as defined in the official Visium HD framework, enabling consistent spatial registration across imaging and sequencing modalities. Sequencing reads were processed using Space Ranger v3 for alignment to the reference genome and assignment to spatial bins on the slide.

**Single-Cell Resolution Inference from 2 µm Visium HD Bins**

Given the highly variable spatial read density, heterogeneous nuclei distribution, and overall low sequencing depth in ccRCC—together with the densely packed tumor microenvironment (TME)—we found that larger bin sizes (8 µm or 16 µm) were suboptimal, as they frequently merged distinct immune and tumor populations. This issue was particularly pronounced due to the small size of lymphocytes relative to tumor clear cells, which introduced substantial ambiguity in downstream deconvolution.

In contrast, direct analysis at the 2 µm bin level was not feasible due to extreme sparsity, likely reflecting limited RNA capture efficiency and cDNA synthesis at this resolution. Moreover, our analytical objectives required relatively pure single-cell–level estimates to distinguish lineage-specific programs, particularly between B-cell and T-cell–associated expression profiles.

To address these constraints, we therefore adopted a single-cell–resolution estimation strategy that integrates histological features, nuclei density, and prior biological knowledge of ccRCC tissue architecture, while avoiding scRNA-seq mapping frameworks such as Tangram that may overfit or impose artificial correspondences between reference cell states and spatial expression patterns. We also evaluated and ultimately moved away from pseudocell-based approaches and benchmarked segmentation tools such as bin2cell and Space Ranger v4 cell segmentation, which are suboptimal in densely infiltrated immune regions. Automated segmentation in Space Ranger was less reliable in this context, whereas H&E morphology provided more informative delineation of cell boundaries in the highly compact immune aggregates characteristic of ccRCC.

Spatial expression data were modeled as a mixture of biological signal and structured technical variation, and a destriping procedure was applied to remove systematic spatial artifacts. Highly abundant transcripts (e.g., IGKC) were excluded to prevent dominance of the low-rank decomposition by a small number of highly expressed genes, and possible contamination of highly expressed antibody producing machinery. Low-confidence genes and ambiguous features were additionally filtered prior to downstream analyses to improve robustness.

Cell-type probabilities were inferred at 2 µm resolution using STHD [5], with reference profiles derived from the Li et al. dataset. These probabilities were then aggregated into 4 µm bins through spatial pooling followed by renormalization, providing a more stable representation for downstream modeling while retaining sub-binning spatial information.

The resulting 4 µm binned data were used as input to a graph-based refinement framework (THOR) [6], which constructs a spatial cell–cell graph based on physical proximity and transcriptional similarity. THOR incorporates nuclei segmentation derived from StarDist applied to aligned H&E images to define cell-aware spatial priors, improving correspondence between transcriptomic bins and underlying cellular boundaries. Within this framework, an anti-shrinking Markov diffusion process propagates information across the graph while preserving sharp transitions between distinct tissue regions.

In our implementation, we further constrained diffusion by pruning edges between high-confidence STHD-assigned cell types that were transcriptionally distinct, preventing over-smoothing across discrete populations while maintaining coherence within homogeneous regions. This constrained diffusion enables recovery of near single-cell–level spatial structure from sparse 2 µm measurements aggregated into 4 µm bins, improving robustness in low-depth, morphologically complex ccRCC tissue while preserving biologically meaningful spatial variation.

### ****Cell type annotation and probabilistic validation in spatial transcriptomics****

STHD-derived annotations were refined using gene set scoring (score_genes, Scanpy) with curated marker sets. Expression was modeled using log-transformed pseudocell estimates derived from probabilistic aggregation of neighboring cells, and scoring was performed directly on these profiles rather than library-size–normalized counts. This approach preserves relative expression structure while stabilizing variance, enabling more robust gene set scoring and spatial comparisons without over-correcting biologically structured signal.

Given the sparsity of spatial transcriptomic data, we benchmarked standard scRNA-seq–style dimensionality reduction and preprocessing pipelines, including log normalization, scaling, PCA, and KNN graph construction, for annotation. These approaches performed sub optimally in this setting, as leading components and neighborhood structure were frequently dominated by technical factors such as library size, sparsity, and local cell density rather than stable biological structure, resulting in unstable variance decomposition across spatial regions. Prior work further suggests that library size in spatial transcriptomics can reflect tissue architecture rather than purely technical noise [7], supporting the use of direct pseudocell-based representations without aggressive normalization or global embedding-based transformations at the annotation stage.

For dimensionality reduction and visualization, we generated latent embeddings using scVI-based models (single-cell Variational Inference; variational autoencoder framework implemented in scVI, scANVI and scVIVA) [8]. These probabilistic models explicitly account for sparsity, dropout, and technical variation by learning a latent representation that separates biological signal from technical noise, making them particularly well suited for spatial datasets characterized by low UMI counts and high dropout rates, where linear methods such as PCA are unstable.

**Histology-Based Spatial Quantification of Tumor and Stromal Architecture**

H&E images were segmented into stromal and tumor compartments using LabKit. Within each region, nuclei were detected and used to compute cell-type proportions as well as nuclei density normalized by stromal area. Lymphocyte surface concentration was defined as the number of lymphocyte nuclei per unit stromal area. Differences between groups were evaluated using two-sided Mann–Whitney U tests.

### ****Spatial distribution of CD8⁺ T cell progenitor–exhaustion transition states****

To characterize the spatial distribution of CD8⁺ T cell states across stromal, tumor, and background histological compartments, we mapped transcriptional states onto segmented tissue regions. CD8⁺ T cell states were modeled along a continuous progenitor-to-exhaustion axis using a progenitor–exhaustion transition score. This score was computed by projecting normalized gene expression signature scores onto a predefined axis spanning progenitor- and exhausted-like programs. Specifically, we calculated the dot product between the normalized score vector and a unit direction vector ([-1, 1]), yielding a single scalar value per cell. Positive values correspond to a more progenitor-like transcriptional program, whereas negative values indicate a more exhausted-like state.

To compare distributions across spatial compartments, we constructed empirical cumulative distribution functions (ECDFs) of transition scores using bootstrap resampling. ECDFs were estimated across resampled cell populations, and curves represent aggregated distributions over iterations to account for sampling variability. For visualization, lower transition scores (reflecting more progenitor-like states) were upweighted to improve resolution of early progenitor populations. Bootstrap-derived confidence intervals were first computed on the unweighted scale and then linearly transformed to the weighted visualization scale. This transformation preserves the relative ordering and overlaps of confidence intervals across groups and does not affect statistical testing, which was performed on the original unweighted distributions. Shaded regions indicate the 5th–95th percentile range of bootstrap ECDFs.

**Spatial Identification, Classification, and Molecular Characterization of Immune Aggregates**

Immune aggregates were defined as spatially coherent clusters of immune cells, motivated by the premise that local cellular composition is a key determinant of immunological function. Candidate aggregates were first identified using HDBSCAN applied to spatial coordinates augmented with scVIVA-derived latent embeddings, enabling joint modeling of physical proximity and transcriptional state. To further enforce spatial coherence, clusters were refined using a k-nearest neighbor graph constructed in physical space, with final aggregates defined as connected components within this graph.

Aggregates were then classified based on the mean variance-stabilized expression of key chemokine markers (CXCL13 and CXCR5) together with the relative composition of stem-like and exhausted CD8⁺ T cell states. This stratification yielded three major classes: lymphoid aggregates, quiescent aggregates, and stimulated aggregates. Lymphoid aggregates were defined by high expression of CXCL13 and CXCR5, consistent with a lymphoid chemokine program. In CXCL13⁻CXCR5⁻ aggregates, classification was instead driven by cellular composition: aggregates enriched for exhausted CD8⁺ T cells were defined as stimulated, whereas those enriched for stem-like CD8⁺ T cells were defined as quiescent. Notably, given the known overlap between effector and exhaustion-associated transcriptional programs in single-cell data, exhaustion-associated signatures may also reflect recent activation states rather than terminal dysfunction.

To identify genes differentially expressed between aggregate classes, we restricted analyses to comparisons with sufficient representation to support robust inference, excluding underpowered groupings such as rare quiescent aggregates in progression contexts and sparsely sampled lymphoid aggregates in the same setting. In non-progression samples, we focused on comparisons between lymphoid and stimulated aggregates.

Given the limited number of slides and substantial biological heterogeneity, conventional pseudobulk differential expression approaches were not suitable. Pseudobulk methods rely on aggregation across samples and assume that observations are independent and identically distributed once summed, an assumption that is often violated in spatial single-cell data. Moreover, they produce integer count matrices that may obscure finer-grained structure present at the single-cell level.

Instead, we used the probabilistic framework of scVI to perform differential expression analysis at single-cell resolution. While scVIVA is an scVI-inspired spatial variational model that incorporates information from neighboring cells to learn spatially aware representations, we opted for scVI because it is more extensively benchmarked and better validated across a wide range of datasets.

A key advantage of scVI is that its training process is focused solely on the gene expression profiles of the cells under analysis, without explicitly incorporating information from other cell types in the surrounding tissue. This reduces the risk of unintended information sharing between distinct cell populations (for example, non-immune cells influencing immune-cell-specific analyses), thereby minimizing potential sources of data leakage in downstream comparisons.

Methodologically, scVI models gene expression within a hierarchical variational framework that explicitly accounts for technical noise, sequencing depth, and batch effects. At the same time, it learns a structured latent representation of biological variation across cells. This probabilistic formulation relaxes the assumption of strict independence between observations required by classical statistical tests, since dependencies and shared structure are captured through latent variables rather than treated as residual noise. As a result, scVI is particularly well suited for spatially organized tissues, where cells are not truly independent but instead exist in correlated microenvironments that reflect underlying biological structure.

Within this framework, individual cells were treated as observations, slide identity was included as a batch covariate, and immune cell type was incorporated as a categorical covariate. Differential expression between aggregate classes was then inferred using the scVI model’s built-in probabilistic DE procedure, which operates on continuous-valued posterior estimates rather than discrete count summaries. These continuous representations (i.e., floating-point-valued latent expressions) are advantageous in this setting, as they reduce sensitivity to sparsity and sampling noise compared to integer-based count aggregation used in pseudobulk approaches such as DESeq2, while retaining uncertainty-aware estimates of gene expression differences.

Genes with the highest posterior probability of differential expression between aggregate classes were subsequently subjected to over-representation analysis using Enrichr, querying the C2 Reactome gene set collection to identify biological pathways associated with distinct immune aggregate states in the ccRCC tumor microenvironment. We prioritized an over-representation framework rather than ranking genes solely by log-fold change, as our objective was comparative characterization of coordinated biological programs rather than classical differential expression focused on effect size magnitude. In this setting, posterior probability provides a more robust measure of reproducible expression differences under the scVI generative model, integrating uncertainty across latent space, batch structure, and cellular heterogeneity. Over-representation analysis is therefore better suited to capture pathway-level structure arising from sets of consistently shifted genes, whereas log-fold change ranking can be disproportionately influenced by sparse or high-variance genes and may not reflect coherent biological programs in complex, spatially structured single-cell data.
